# DIP-MS: A novel ultra-deep interaction proteomics for the deconvolution of protein complexes

**DOI:** 10.1101/2023.03.22.533843

**Authors:** Fabian Frommelt, Andrea Fossati, Federico Uliana, Fabian Wendt, Xue Peng, Moritz Heusel, Bernd Wollscheid, Ruedi Aebersold, Rodolfo Ciuffa, Matthias Gstaiger

## Abstract

Most, if not all, proteins are organized in macromolecular assemblies, which represent key functional units regulating and catalyzing the majority of cellular processes in health and disease. Ever-advancing analytical capabilities promise to pinpoint lesions in proteome modularity driving disease phenotypes. Affinity purification of the protein of interest combined with LC-MS/MS (AP-MS) represents the method of choice to identify interacting proteins. The composition of complex isoforms concurrently present in the AP sample can however not be resolved from a single AP-MS experiment but requires computational inference from multiple time-and resource-intensive reciprocal AP-MS experiments.

In this study we introduce Deep Interactome Profiling by Mass Spectrometry (DIP-MS) which combines affinity enrichment with BN-PAGE separation, DIA mass spectrometry and deep-learning-based signal processing to resolve complex isoforms sharing the same bait protein in a single experiment.

We applied DIP-MS to probe the organisation of the human prefoldin (PFD) family of complexes, resolving distinct PFD holo- and sub-complex variants, complex-complex interactions and complex isoforms with new subunits that were experimentally validated. Our results demonstrate that DIP-MS can reveal proteome modularity at unprecedented depth and resolution and thus represents a critical steppingstone to relate a proteome state to phenotype in both healthy and diseased conditions.

## Introduction

Understanding how different proteins are spatially organized into functional modules catalyzing and controlling numerous biochemical and cellular processes underlying distinct phenotypes is one of the main goals of molecular systems biology. Protein complexes, defined here as stable assemblies that can be isolated by biochemical means, are key regulators and catalysts of cellular functions. Far from being invariant assemblies, protein complexes are contextual and have been shown to adapt to the cellular type or state by changing their subunit composition, stoichiometry, localization and abundance of expression [1, 2]. Gaining insights into the nature of protein complexes in the context of a particular cellular type or state, is therefore essential to understand the biochemical processes underlying genotype-phenotype relationships [3]. Affinity purification mass spectrometry (AP-MS) [4, 5] has been the method of choice for the analysis of protein complexes due to its high sensitivity via enrichment of the bait protein and its interaction partners. However, AP-MS of a single bait identifies direct as well as indirect interactors that may not belong to the same complex but rather be part of different complexes concurrently present in the AP sample. Therefore, PPI data from several reciprocal AP-MS experiments using the interactors as baits are typically needed to deconvolve the MS data into distinct molecular entities [6].

As an alternative to AP-MS protein correlation methods exemplified by size exclusion chromatography coupled to mass spectrometry (SEC-MS) [7, 8] and blue native-PAGE (BN-PAGE) [9] coupled to MS [10] have been introduced. They fractionate native complexes in mild cell extracts by their physical properties (hydrodynamic radius and size, respectively) under near-native conditions and each fraction is subsequently profiled by MS. The resulting protein profiles across the fractionation dimension are then used as proxies to define protein complexes within the sample. While these approaches can identify multiple concurrent protein complexes involving overlapping proteins due to the additional separation in the molecular weight dimension, the information is typically limited by the sensitivity range of the analytical instrument, the total sample loading capacity of the column and the limited resolution of the SEC columns [11, 12]. Collectively, these factors limit the general utility of co-fractionation-MS methods for detecting (i) complexes present at low abundance, (ii) complex components present in substoichiometric amounts, and (iii) the resolution of different complex instances containing the same core subunits but different accessory proteins.

Here we introduce Deep Interactome Profiling by Mass Spectrometry (DIP-MS), a method combining affinity purification with native-fractionation by BN-PAGE and data independent acquisition mass spectrometry (DIA-MS) and a bespoke data analysis software package. DIP-MS combines the capacity of AP-MS to enrich the interactome of a target protein with the ability of native fractionation-MS approaches to resolve different complexes, and complex instances sharing the same target protein. BN-PAGE was chosen both because it does require significantly lower amounts of samples compared to SEC, and because it is well suited for multiplexing and comparison across conditions within a single PAGE gel. In addition to introducing improvements in the experimental protocol to increase throughput and reproducibility, we developed *PPIprophet*, a data-driven neural network-based protein complex deconvolution system.

Compared to the few prior studies that combined AP with fractionation MS [13–15], DIP-MS provides three critical improvements: (i) a novel miniaturized sample preparation procedure in a filter plate format that requires 10 times less material than traditional chromatography-based separation [16, 17] and achieves high reproducibility; (ii) a DIA-MS acquisition scheme coupled with a 24 minutes injection to injection gradient chromatography system that allows for measuring of up to 60 samples per day, thus increasing drastically sample throughput compared to prior methods; (iii) a deep-learning based framework trained on more than 1.5 million binary interactions from 33 SEC, IEX and BNP datasets, which enabled prediction of protein-protein interactions (PPIs), identification of multiple instances of protein complexes, and ultimately the reliable deconvolution of complex profiling data into functional modules.

To benchmark and explore the potential of DIP-MS for large scale PPI profiling, we analyzed the interactome of the human prefoldin proteins. Prefoldins play a central role in the cellular proteostasis network because they stabilize nascent proteins in interplay with other chaperones [18–20]. They are best known as part of the evolutionarily conserved heterohexameric canonical prefoldin (PFD) complex, a cytosolic ∼120 kDa ATP-independent chaperone comprising two different ∼23 kDa *α*- and four different ∼14 kDa *β*-subunits [18, 21]. In addition to the prototypical PFD complex, complexes containing prefoldin subunits have been implicated in a range of cellular processes including neurodegeneration [22–24] and degradation of misfolded proteins [25] and detected in different cellular compartments including the cytosol, mitochondria and nucleus [26, 27]. Further, prefoldin and prefoldin-like proteins form the prefoldin-like module (PFDL) [28]; they can assemble in supercomplexes, such as the chaperonin CCT/TRiC-PFD [29] and most prominently the PAQosome (also known as R2TP/PFDL or URI complex), a HSP90 chaperone complex, which assists in mammalian cells in assembly and maturation of large RNA-binding protein assemblies [30–34] (see Table 1 and Extended data Fig. 1).

**Fig. 1.**
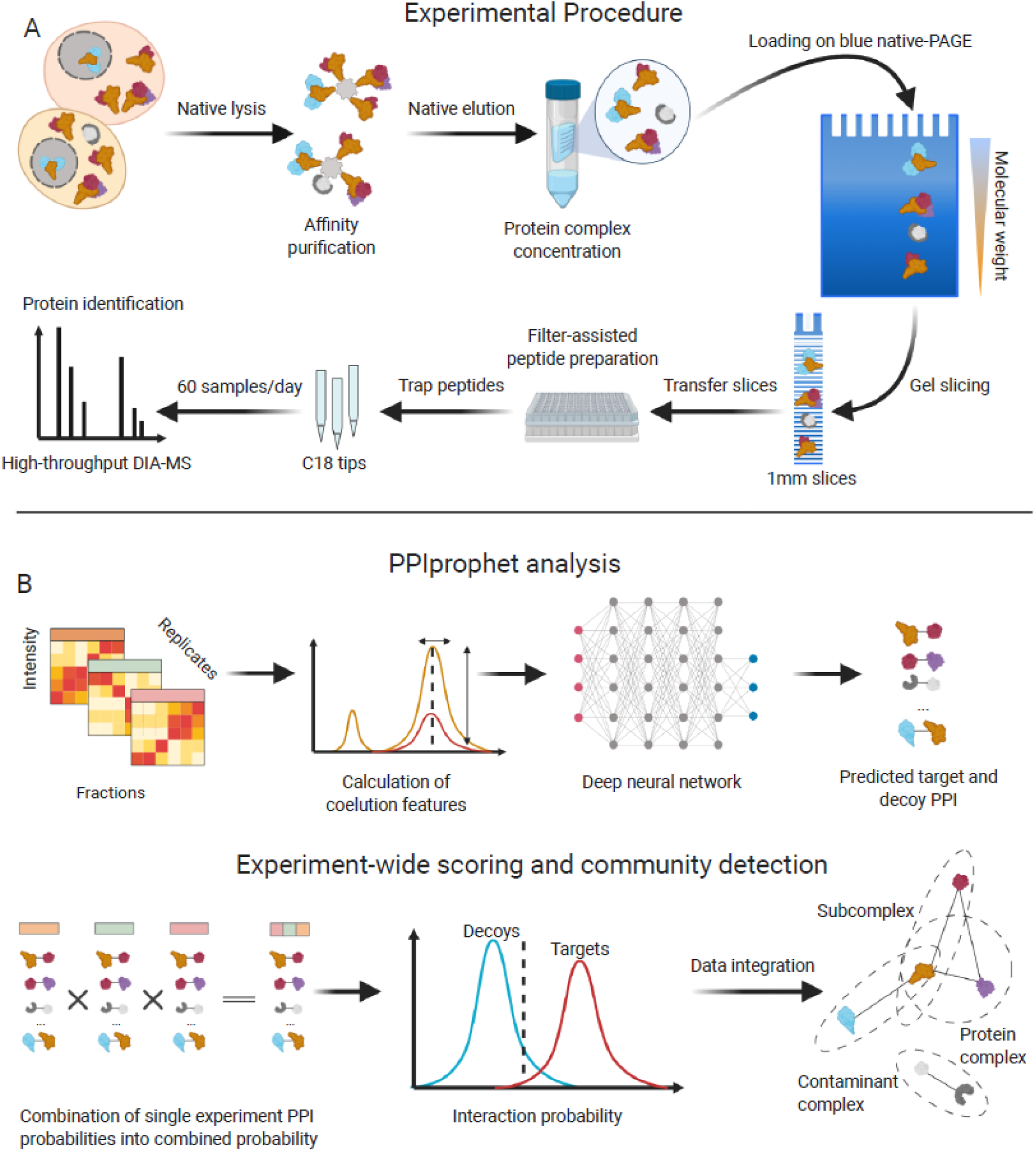
Schematic of experimental and computational DIP-MS workflow. **A**. Experimental workflow, including sample preparation, affinity purification, gel-based fractionation and DIA acquisition. **B.** Computational framework encompassing first, generation of all possible PPIs in the data and then prediction of PPI using deep learning. Interaction probabilities are then combined and further filtered using a target-decoy competition approach. Complexes are derived from the resulting interaction network by integration with available databases or by using a data-driven approach.

**Table 1.** Core-subunits in the prefoldin network.

| Gene Name | UniProtKB accession | Complex | Bait in reciprocal AP-MS | References |
| --- | --- | --- | --- | --- |
| PFDN1 | O60925 | Canonical prefoldin | Bait | [18-21, 29] |
| PFDN2 | Q9UHV9 | Canonical prefoldin | Bait | [18-21, 29] |
| VBP1 | P61758 | Canonical prefoldin | Bait | [18-21, 29] |
| PFDN4 | Q9NQP4 | Canonical prefoldin | Bait | [18-21, 29] |
| PFDN5 | Q99471 | Canonical prefoldin | Bait | [18-21, 29] |
| PFDN6 | O15212 | Canonical prefoldin |  | [18-21, 29] |
| ASDURF | L0R819 | Prefoldin-like/URI |  | [52] |
| PDRG1 | Q9NUG6 | Prefoldin-like/URI | Bait | [28, 31, 34, 52] |
| PFDN2 | Q9UHV9 | Prefoldin-like/URI | Bait | [28, 31, 34, 52] |
| PFDN6 | O15212 | Prefoldin-like/URI |  | [28, 31, 34, 52] |
| URI1 | O94763 | Prefoldin-like/URI |  | [28, 31, 34, 52] |
| UXT | Q9UBK9 | Prefoldin-like/URI | Bait | [28, 31, 34, 52] |
| RUVBL1 | Q9Y265 | R2TP | Bait | [28, 31, 34, 52] |
| RUVBL2 | Q9Y230 | R2TP | Bait | [28, 31, 34, 52] |
| RPAP3 | Q9H6T3 | R2TP | Bait | [28, 31, 34, 52] |
| PIH1D1 | Q9NWS0 | R2TP | Bait | [28, 31, 34, 52] |
| WDR92 | Q96MX6 | PAQosome<br>associated protein |  | [28, 31, 34, 52] |
| POLR2E | P19388 | PAQosome<br>associated protein |  | [28, 31, 34, 52] |

To gain further insights into the landscape of prefoldin complexes we performed DIP-MS on the PDF and PFDL complexes, respectively, using PFDN2 and UXT as bait proteins, whereby PFDN2 is shared between different complex instances and UXT is exclusive for PDFL. Analysis with *PPIprophet* and comparison of the results with those obtained by AP-

MS and SEC-SWATH identified most known prefoldin complexes in a single experiment and identified 319 PFDN2-specific interactors, thus substantially outperforming the state-of the art approaches. DIP-MS not only recapitulated the composition of all reported PFD complexes but also quantified their stoichiometry and additionally suggested the existence of stable sub-assemblies and supercomplexes. Further it supported the identification of different instances of the respective complexes, revealed a previously unknown PFD homolog and deconvolved the PFD and PFDL-complex landscape into multiple complex instances. In summary, we introduce a new method to quantitatively study the organization of the proteome at unprecedented resolution and sensitivity.

## Results

### Overview of the DIP-MS Method

To identify the complexes contained in an affinity purified sample we developed and implemented deep interactome profiling (DIP-MS), a workflow combining affinity purification with size-based fractionation of enriched complexes by blue native-PAGE (BN-PAGE) separation, fast DIA-MS and a dedicated software tool.

The steps of the DIP-MS experimental workflow are illustrated in (Fig. 1A). The protein complexes containing the bait protein were purified using an affinity tag and subjected to BN-PAGE to separate complexes by their apparent molecular weight (MW). After recording the stained gel pattern, the gel was cut into 70 slices of 1 mm width, which were then individually processed. To this end, we devised a fast and reproducible filter aided in-gel digestion preparation protocol in a 96-well plate format. Finally, proteolyzed peptides from each fraction were measured with quantitative DIA-MS [35] coupled with a short LC gradient [36] for each consecutive fraction. The resulting comigration (from here on coelution) matrices of peptide fragment ion spectra (Fig. 1B) were processed via *PPIprophet* to infer quantitative protein electrophoretic elution patterns, assembly states, and protein-protein interactions (PPIs) Protein profiles showing multiple peaks were further deconvolved to infer multiple assemblies, thereby identifying in a single DIP-MS experiment multiple complexes and sub-assemblies containing the bait protein. This greatly enhanced the coverage and resolution of the interaction network.

### *PPIprophet*: a deep-learning powered framework for PPI prediction and complex inference

The *PPIprophet* software was specifically developed to analyze results generated by a DIP-MS experiment. The software predicts PPIs from affinity-purified, fractionated samples. Specifically, the following information is provided by DIP-MS datasets: i) identification of protein complexes containing the bait protein, ii) subunit stoichiometry and approximate molecular weight based on the electrophoretic mobility of separated complexes, and iii) co-fractionation of prey proteins implying prey-prey interactions which are usually invisible to AP-MS without extensive reciprocal purification.

In developing *PPIprophet*, we trained a deep neural network model (see method for details) for PPI prediction using more than 1.5 million unique PPIs extracted from databases containing data from different types of co-fractionation measurements and organisms [37]. By utilizing STRING and BioPlex as ground truth, this DNN model achieved outstanding performance with a ROC of 0.995 on our independent test set of 335’071 PPIs (Table 2 and method for details on model training and testing datasets), showing the flexibility of deep learning for this task compared to previously employed correlation-based approaches [13]. To reduce the false discovery rate (FDR) due to spuriously comigrating proteins, *PPIprophet* performs FDR control using data-generated decoy PPIs. By exhaustively mapping and predicting all possible PPIs represented by the data, the software tool generates a weighted network (Fig. 1B) which is then used for further protein complex identification. To distinguish complex components from contaminants (a notorious challenge for traditional AP-MS scoring algorithms), we devised a interaction metric (W-score, a customized WD-score adapted from CompPASS [38]) that employs specificity and selectivity to filter copurifying proteins, by performing *in silico* affinity purifications (see method for details). Finally, for identification of protein complexes, *PPIprophet* can be used either in a hypothesis testing mode where the PPI network is deconvolved into complexes by superimposition of available complex knowledge or in an entirely data-driven mode using MCL clustering [39].

**Table 2.** Prediction performance of PPIprophet.

| Metric | Value |
| --- | --- |
| Accuracy | 0.984 |
| Precision | 0.969 |
| Recall | 0.9545 |
| F1 score | 0.961 |
| ROC AUC | 0.995 |

### Benchmarking of DIP-MS against AP-MS and SEC-MS workflows for interactome analysis

To evaluate the performance of the method in relation to the present standard methods, we performed DIP-MS using complexes affinity isolated with prefoldin subunit 2 (PFDN2) as bait protein. Briefly, we purified ectopically expressed, SH-tagged PFDN2 from 6 x 10^8 HEK293 cells and separated the eluate by BNPAGE in biological triplicates (Supplementary Fig. S1A-F). Using a set of isotopically labeled spike-in peptides with the tag sequence, we estimated the sample load at approximately ∼2 *µ*g of purified bait-complex per lane (Supplementary Data S1). Gel lanes were then sliced into fractions, and the proteolyzed content of each fraction was analyzed in DIA-MS mode using a short 20-minute gradient (Methods) and analyzed the data with *PPIprophet*. Overall, we could reconstruct the migration profile of 1465 proteins (Supplementary Data S2). Next, to evaluate the performance of the method, we defined a ground truth benchmark which is comprised of two distinct interaction lists. The first is a literature curated list of 16 well characterized PFD and PFDL core complex subunits (Supplementary Table S1). The second consists of the combination of the first list together with the binary protein interactors of all subunits of the PFD and PFDL complexes, the PFDL containing PAQosome and CCT/TRiC reported in BioPlex [5], BioGRID [40] and IID [41].

Lists I and II are based on more than 556 studies and include a total of 1891 interactors. We compared the DIP-MS results with data generated by two orthogonal methods: (i) SEC-MS, analyzed with CC-Profiler [16]; and (ii) an in-house generated reciprocal AP-MS dataset using 11 core-interactors from the PFD/PFDL interaction network as bait proteins (Supplementary Table S2) analyzed with SAINTexpress [42]. The AP-MS dataset recovered 77 reported PFD/PFDL interaction partners, including the components of the two core complexes (∼7.5% of the curated list, Fig. 2A). The SEC-MS dataset identified 151 proteins, (14% of the curated list). In the SEC-MS dataset the PFD complex was identified from the profiles of coeluting proteins, whereas the PFDL complex could not be identified compared to both DIP-MS experiments (Supplementary Fig. S2A). Comparison to 36 coelution studies of human cell extracts showed that on average 3 out of 6 PFDL subunits were recovered with only 2 studies recovering all 6 subunits (Supplementary Fig. S2B). Remarkably, the DIP-MS dataset identified 319 PFD/PFDL interaction partners in total, 187 more than both SEC-MS and AP-MS, recovering ∼30% of the interactors in public databases (Fig. 2A).

**Fig. 2.**
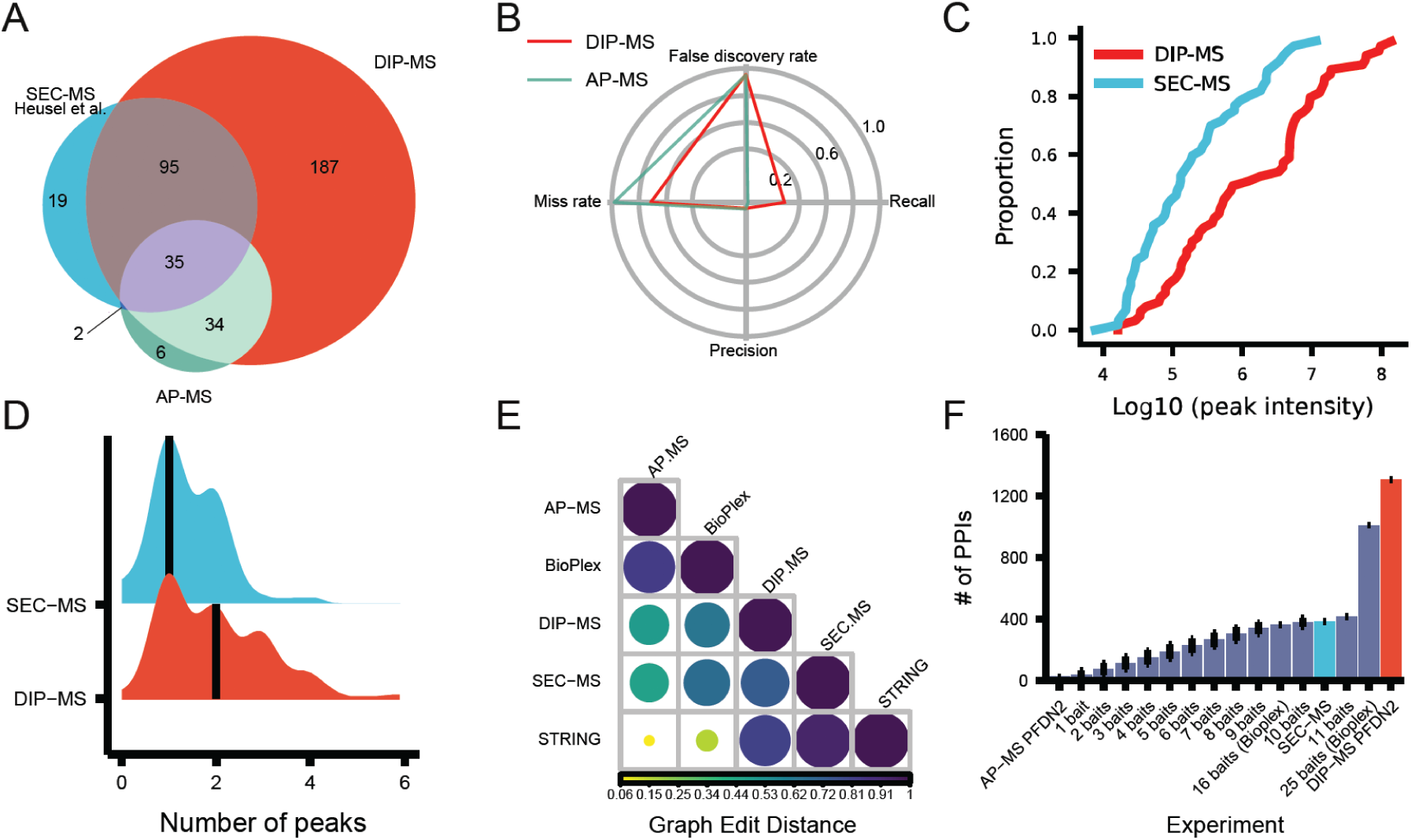
Benchmark of DIP-MS versus other techniques for interactome analysis. **A**. Venn plot showing the overlap in identified PFDN2 interactors in SEC-MS, AP-MS (across 11 core-subunits of PFD and PAQosome) and DIP-MS from the curated list of known PFDN2 interactors. **B.** Radarplot showing the performance of DIP-MS over AP-MS. **C.** Empirical Cumulative Distribution Function (ECDF) plot for PFD/PFDL associated proteins identified in the PFDN2 DIP-MS experiment (average across replicates) and the SEC-MS from Heusel et al. [16]. X-axis represents the log10 MS2 abundance integrated over a protein coelution profile in the SEC-MS dimension, defined here as a protein peak. Y axis represents the proportion of proteins peaks at a specific abundance. **D.** Number of coelution peaks per PFD/PFDL associated proteins, identified in both, SEC-MS and DIP-MS. Solid black line represents the median for SEC-MS (1) and for DIP-MS (2) while Y axis represents the kernel density for the SEC-MS (blue density) and the DIP-MS (red density). **E.** Graph Edit Distance (GED) matrix between different techniques and databases. Color code represents the GED similarity from highly dissimilar (yellow, small circles) to isomorph graphs (dark purple, large circles). **F.** Barplot showing the number interactions recovered by different methods. Bars represents standard deviation with n = the total number of combinations possible from the correspondent bait number. Different techniques are highlighted by different bar colors.

Importantly, higher coverage benchmark data did not significantly affect error rates, as we observed greater recall and similar precision compared to our in-house generated AP-MS data using the manually curated set as ground truth, suggesting that DIP-MS generates much larger and denser interaction networks at no cost of precision (Fig. 2B).

We hypothesized that DIP-MS can recall a larger fraction of proteins in the benchmarking dataset compared to global co-fractionation MS because of an increased depth of protein identification provided by the initial bait-enrichment step. To test this hypothesis, we plotted the intensity of the MS signals of the benchmark proteins as surrogates of protein abundance in the DIP-MS and SEC-MS datasets (Fig. 2C). We found that the signal intensities in the SEC-MS data for all the proteins in the target set covered a range of ∼3 logs of abundance. In contrast, signal intensities in the DIP-MS data covered a dynamic range of ∼4 logs, suggesting increased coverage of low-abundance proteins from the target set. It is important to note that the SEC-MS experiment used as benchmark was measured in a different MS system compared to our DIP-MS so lower or higher absolute abundance should not be considered as proxy of coverage, while the proportion of signal (i.e., Y axis) in the ECDF plot is magnitude-agnostic allowing comparison between different instruments.

Because the enrichment step lowers the detection threshold for low-abundant proteins, and, at the same time, significantly reduces the complexity of the sample in each gel fraction compared to the unseparated sample analyzed in SEC-MS, we hypothesized that DIP-MS should also be able to resolve a greater number of complexes than SEC-MS. To test this, we applied the same signal processing and peak-picking algorithm to all benchmark proteins identified by both methods (n=123) and calculated the number of distinct peaks detected for each protein. In this context, we operationally defined a protein peak as a signal having minimal width of 3 fractions and minimal height above background of 0.2 on a 0-1 signal scale. Strikingly, when comparing the distributions derived from SEC-MS and DIP-MS, the latter showed significantly increased number of peaks (1 vs 2, p=0.00123) (Fig. 2D), in line with our expectation of increased elution signals detectability due to extended dynamic range and interactome enrichment. Overall, these analyses support the notion that the combination of affinity-enrichment and fractionation boosts the depth, resolution, and, therefore, overall performance, of the method.

Next, we asked whether PPI networks generated by DIP-MS have different properties (topology and connectivity) compared to those generated by SEC-MS and AP-MS. To determine which method most closely recapitulated the network topology of a large-scale PPI database, we calculated the Graph Edit Distance (GED) between the subnetworks encompassing all the 475 PFD/PFDL proteins from the target list identified in all three experiments (intersection of quantified proteins across the DIP-MS, SEC-MS and reciprocal AP-MS with 11 baits experiments) and two representative PPI databases (STRING and BioPlex). These 475 proteins represent only the proteins identified, not necessarily predicted as positive interaction partners, hence offer an unbiased metric of algorithm performance in network reconstruction. GED is a measure of topological similarity between networks, taking the value of 1 for identical graphs. Lower values indicate increasingly diverging networks (Materials and Methods for details). As expected, we observed high GED for datasets generated by the same technique. Specifically, the AP-MS derived network was closer to the large scale AP-MS dataset BioPlex than SEC-MS and DIP-MS. The latter networks were more similar to STRING (0.94 and 0.91, respectively), whereas the AP-MS derived network was vastly different from STRING (GED score of 0.06) as shown in Fig. 2E. This indicates that DIP-MS recapitulates the topology of the graph as well as SEC-MS, but, critically, does not rely on prior knowledge and therefore allows discovery of new unreported complexes. Finally, we compared the number of PPIs, a direct proxy for the density of the network, generated by DIP-MS, SEC-MS and AP-MS. A single-bait DIP-MS experiment (PFDN2) was sufficient to generate a network with 1306 PPIs. Strikingly, a subset of the SEC-MS data containing the same proteins identified in DIP-MS from the target list led to a significantly less-well connected network of 386 PPIs. To compare DIP-MS and AP-MS data derived networks, we asked how many AP-MS experiments would be required to reconstruct a network as dense as the one generated by one DIP-MS measurement. To this end, we queried the BioPlex interaction network in HEK293 cells and selected all interactions encompassing either 16 baits (11 core PFDN and PFDL subunits and 5 proteins from the R2TP module) or 25 baits (11 PFD/PFDL-core, 5 from the R2TP module and 9 from CCT/TRiC). We found that even using 25 baits from BioPlex (core subunits + R2TP + CCT/TRiC) yielded approx. 20% fewer PPIs than the PPIs retrieved by a single DIP-MS experiment (1011 vs. 1306) (see Fig. 2F).

In summary, our benchmarking data indicates that DIP-MS data, when compared to SEC-MS data of whole cell extracts and to AP-MS data using reciprocal tagging schemes, has a broader dynamic range, captures the separation behavior of proteins at higher resolution, and generates more extensive and denser networks and recapitulated a significantly larger portion of the ground truth.

### Global organization of Prefoldin and Prefoldin like complexes

We next applied DIP-MS to generate a detailed map of prefoldin and prefoldin-like complexes using the alpha prefoldin subunit UXT and the beta prefoldin subunit PFDN2 as baits. PFDN2 is a shared subunit of the PFD, PFDL and PAQosome multimers, whereas UXT was reported as subunit for the PFDL and PAQosome complexes (Extended data Fig. 1) [28, 43]. Similarly, to the PFDN2 DIP-MS experiment above, we purified SH-tagged UXT from 6 x 10^8 HEK293 cells in biological triplicates, separated the eluate on a native-PAGE gel (Supplementary Fig. S3A-F) and after initial data processing quantified the elution profiles for 737 proteins (Supplementary Data S2).

Across the two DIP-MS experiments we profiled 1’513 proteins after applying an initial data processing module of the R-package CCprofiler [16] for sibling peptide correlation-based filtering and conversion of peptide-level features into protein-level features. Using *PPIprophet* to analyze the protein profiles we detected 6’762 (PFDN2) and 5’682 (UXT) high-confidence (combined probability across replicates >= 0.9) binary interactions, resulting in a network containing a total of 11’552 unique interactions and 939 proteins. We first asked what portion of the already characterized prefoldin and prefoldin-like complexes were recalled from this data. In our combined DIP-MS derived network, we identified all of the previously reported PFD/PFDL assemblies (Extended Fig. 2) such as PFD, PFDL and the PFDL containing PAQosome [18, 28]. Further we identified coelution groups comprised of the R2TP proteins RUVBL1/2 and the R2TP adaptors/regulatory subunits RPAP3 and PIH1D1 and the CCT/TRiC-PFD complex.

The PFD and PFDL interacting with the URI1 binding protein POLR2E, (Fig. 3A) complexes were base-peak resolved and could be readily identified by naïve clustering procedures (Supplementary Fig. S4A). Besides their coelution at the expected MW, these two complexes additionally migrated at a high MW, where we found that PFD and PFDL partake in two distinct supercomplexes. The first is a PFD-CCT/TRiC supercomplex (Extended data Fig. 2 coelution group 6), and the second is the PAQosome (Extended data Fig. 2 coelution group 7) which comprises the fully assembled R2TP and PDFL. Remarkably, our data captured the reported variant of RUVBL1/2 as a hetero-dodecameric complex (PDB id: 2XSZ) [44–46] lacking the adaptor/regulatory proteins RPAP3 and PIH1D1 (Fig. 3B) which are part of the R2TP-core. It was proposed that RUVBLs cycle between a double and single ring form which may give rise to specialized chaperone function of the R2TP core [47]. Surprisingly, for the two adaptor proteins PIH1D1 and RPAP3 [48], we identified three separate peaks which suggests the presence of multimeric sub-assemblies formed by RPAP3 and PIH1D1. We also identified the two R2TP subunits at the PAQosome coelution group. Indeed, in a recently published R2TP structure, RPAP3 binds to PIH1D1, and this assembly is recruited to the RUVBL1/2 hexamer by binding of the C-terminal domain of RPAP3 to the RUVBLs [47, 49, 50]. Further, in recent work, the structure of a RPAP3:PIH1D1 complex was resolved [50], which confirms our observation of an independent RPAP3:PIH1D1 subassembly (Supplementary Data S3). In prior work RPAP3:PIH1D1 showed high affinity binding to HSP90, and indeed in the lower MW peak group of the adaptor peak we identified coelution with HSP90 complex subunits (Supplementary Fig. S4B).

**Fig. 3.**
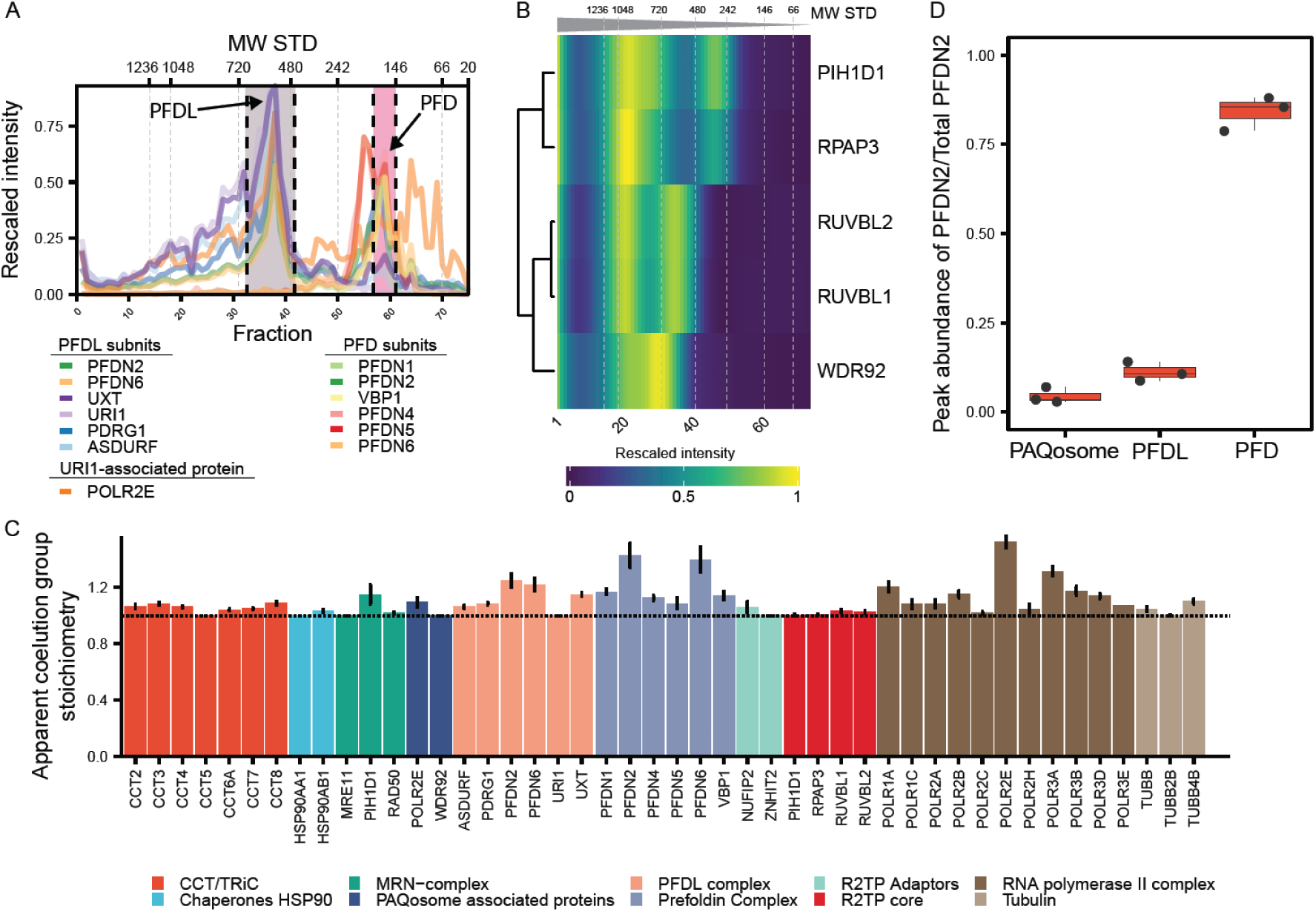
Identification of assemblies from PFDN2 and UXT DIP-MS experiments. **A**. Coelution profiles PFD and PFDL complexes in the PFDN2 DIP-MS experiment. X axis represents the gel slice number, while the Y axis shows the rescaled protein intensities (sum of TOP2 peptides, MS2 based protein intensity, summed up for each protein across all DIP-MS experiments before scaling). Region between the black dotted lines delimits the peak group selected as assembled PFD and PFDL. Molecular weights of the standards are reported as additional X axis on top. **B.** Heatmap of the R2TP/PAQosome subunits Ruvbl1/2, RPAP3 and PIH1D1 after Ward’s distance-based clustering for the UXT-DIP-MS purification. Cell values represents the rescaled intensity averaged across the triplicates.**C.** Barplot showing the stoichiometry obtained from the DIP-MS experiments of UXT and PFDN2 for representative complexes in the PFD and PFDL interaction network as defined in Supplementary Table S1. Dotted line highlights a stoichiometry of 1.00, which correspond to the literature-reported value for PFDN, PFDL as well as CCT/TRiC. Error bars indicate the standard deviation across 6 DIP-MS experiments (PFDN2, n=3; and UXT, n=3). **D.** Boxplot showing the ratio of PFDN2 (bait) signal in the PFD, PFDL or PAQosome complex over the total PFDN2 signal. Box shows the interquantile range (IQR) and the whiskers 1.5 X IQR. Black line represents the median while each dot shows one independent replicate (n=3).

Importantly, of the 7 distinct assemblies identified in our DIP-MS experiment only the highly abundant prefoldin complex was identified and resolved in previous SEC-MS experiments [7]. Of note, PFDL and PAQosome subunits were identified in conventional whole cell lysate SEC-MS, but did not show detectable coelution (i.e overlapping peaks at high molecular weight) [16], exemplifying the increased sensitivity and scope for discovery of low abundant protein complexes using the DIP-MS technology.

Next, we asked whether DIP-MS could provide reliable quantification of the identified assemblies, focusing on stoichiometry and occupancy, defined as the fraction of the total bait protein detected that is associated with a specific interactor or complex. To achieve this, we first calculated the apparent subunit stoichiometry of the complexes (defined here as the ratio between the protein having the lowest full-width half max peak area in a complex and all the other proteins in the same peak group) and compared it to the reported stoichiometries from structural studies (Fig. 3C). Complexes known to have a 1:1 subunit stoichiometry such as PFD (PFDN1, PFDN2, VBP1, PFDN4, PFDN5, PFDN6), PFDL (PFDN2, PDFN6, UXT, URI1, PDRG1, ASDURF), CCT/TRiC (TPC1, CCT2, CCT3, CCT4, CCT5, CCT6A, CCT6B, CCT7, CCT8), RNA polymerase (RNAPs), R2TP (RUVBL1, RVUBL2, RPAP3, PIH1D1) were indeed close to their reported stoichiometry even in the case of the PFD complex and the PFDL which ectopically express affinity tagged subunits PFDN2 and UXT, respectively. These results indicate that DIP-MS derived stoichiometry values are in agreement with those derived by structural biology methods and, more broadly, that complex stoichiometry tend to be maintained regardless of the level of the cellular expression of individual subunits as previously proposed [1]. Next, we calculated the occupancy for both PFDN2 and UXT. Our data suggests that the PFD complex dominates the PFDN2 interaction network. It accounts for ≈ 90% of the total PFDN2 signal, while only a fractional amount (≈ 8%) of PFDN2 can be found in the PFDL complex (Fig. 3D) and less than 1% of the PFDN2 signal is in the PAQosome. In contrast, when we performed DIP-MS using the PFDL subunit UXT as bait (Supplementary Fig. S3A-F), we enriched the PFDL peak compared to the PFDL peak within the PFDN2 DIP-MS experiment by more than 5 fold (≈ 75% of the total PFDN2) and the PAQosome peak by more than 20 fold, while simultaneously almost completely depleting the PFD peak, which is a likely contaminant in the UXT purification (Extended data Fig. 3A). When comparing the intensity for the same complex between the two tested bait proteins we found significant enrichment of PFDL in UXT (Log2FC 1.88) and depletion of the PFD complex (Log2FC -12.4) as shown in Extended data Fig. 3B. For the PAQosome we did not observe significant enrichment (Log2FC 0.016) between the two baits. DIP-MS with UXT thus enabled the validation of the PAQosome supercomplex which was enriched to a similar extend in the DIP-MS of PFDN2.

Comparison of our results with previously published SEC-MS data [16] indicates that, as expected, the low expression of PAQosome and PFDL (average of 69 normalized transcripts per million (nTPM) vs. 203 nTPM for exclusive subunits) results in them having only a fractional amount of the total signal (3.3e5 for PFDL and 2.5e5 for the PAQosome in SEC-MS compared to 1.0e7 and 6.7e6 in UXT DIP-MS, see Extended data Fig. 3C-E and Supplementary Data S4). This, in turn results in poorer detection of coelution, a problem substantially alleviated by the pre-fractionation enrichment in DIP-MS.

Finally, to further highlight the versatility of the method, we performed absolute quantification of proteins as previously described [51]. Briefly, an external calibration curve was built using a synthetic heavy peptide which correspond to the tryptic peptide of the affinity tag. Based on this calibration curve, we estimated the absolute amount of PFDN2 in the DIP-MS inputs prior to separation being ≈ 2.24 µg (see Supplementary Data S1). Consequently, the PFD peak contains ≈ 2.02 µg of PFDN2, while ≈ 22.4 ng are present in the PFDL. The lowest signal measured for the PFDN2 across the DIP-MS gradient was estimated at ∼22 fmol (average of DIP-MS PFDN2 replicate 1 and 2).

Overall, these two DIP-MS experiments provided the thus-far most exhaustive account of the organization and architecture (i.e., stoichiometry) of prefoldin complexes. In addition to recalling prior knowledge, we were able, from the same data, to identify two structural variants of PAQosome subunits: the reported RUVBL1/2 heterohexamer and the RPAP3:PIH1D1 subassembly. The absolute quantification of the DIP-MS input allowed us to quantify that around 1 – 3 μg of purified bait is sufficient to resolve bait-containing protein complexes employing DIP-MS.

### Discovery of an alternative PDRG1-containing PFD complex

Having shown that the DIP-MS method can capture the two major assemblies (PFD and PFDL) in the prefoldin network and their repartition in super complexes (CCT/PFD and the PAQosome), we turned our attention to proteins that were predicted by *PPIprophet* as genuine complex components but that were (i) previously not known to be part of PFD or PFDL complexes or (ii) whose role has not been extensively clarified.

To achieve this, we took the combined results from *PPIprophet* analysis of the UXT and PFDN2 DIP-MS experiments and selected all subunits of the PFDL containing PAQosome. Overall, we assigned the 16-core subunits of the PAQosome to multiple subassemblies in our DIP protein-coelution profiles, which allowed us further to reveal the underlying molecular architecture of this interaction network (see Extended data Fig. 2 and Supplementary Data S3). Specifically, we focused on two proteins, PDRG1 and ASDURF which were previously reported as PFDL or PAQosome subunits [52].

The PFDN2 DIP-MS data showed that PDRG1 eluted in three separate peaks compared to the UXT-DIP-MS for which we only obtained two peaks (Extended data Fig. 4A). The first two MW peaks corresponded to the previously reported PFDL complex and the PFDL containing PAQosome. Additionally, we discovered that, in the third and lowest MW peak, PDRG1 coeluted with canonical PFD subunits as shown in Figure 4A. This finding is further supported by results from clustering the interaction probabilities from *PPIprophet* within the PFDN2 DIP-MS experiments of PDRG1 with PFD subunits and PFDL (Fig. 4B). Interestingly, the non-canonical *β*-PFD subunit PDRG1 was originally termed PFDN4r related protein due to its sequence homology (30% identity) to the canonical *β*-PFD subunit PFDN4 [28] (Extended data Fig. 5A, Supplementary Data S5 and S6). To further validate our finding, we performed reciprocal AP-MS of four PFD subunits reported to be mutually exclusive for the canonical complex (PFDN1, VBP1, PFDN4, PFDN5) and PDRG1 (Fig. 4C; Supplementary Data S7 contains results for all reciprocal AP-MS conducted within this study, see methods regarding scoring and filtering strategy). We recovered PDRG1 as a high confidence interactor of all four baits used, thereby confirming the interaction with canonical PFD-subunits. At the same time, the AP-MS results using PDRG1 as bait protein identified both canonical PFD subunits and PFDL and PFDL containing PAQosome complex members as interactors, consistent with the DIP-MS results (Fig. 4B). Interestingly, when comparing the PFD stoichiometry across the various reciprocal control AP-MS experiments, after normalization for the PFD subunit VBP1, we observed a significantly lower stoichiometry for PFDN4 in the PDRG1 purification compared to all other purifications, including the control samples (Extended data Fig. 5B) suggesting that PFDN4 is a contaminant in the PDRG1 purification.

**Fig. 4.**
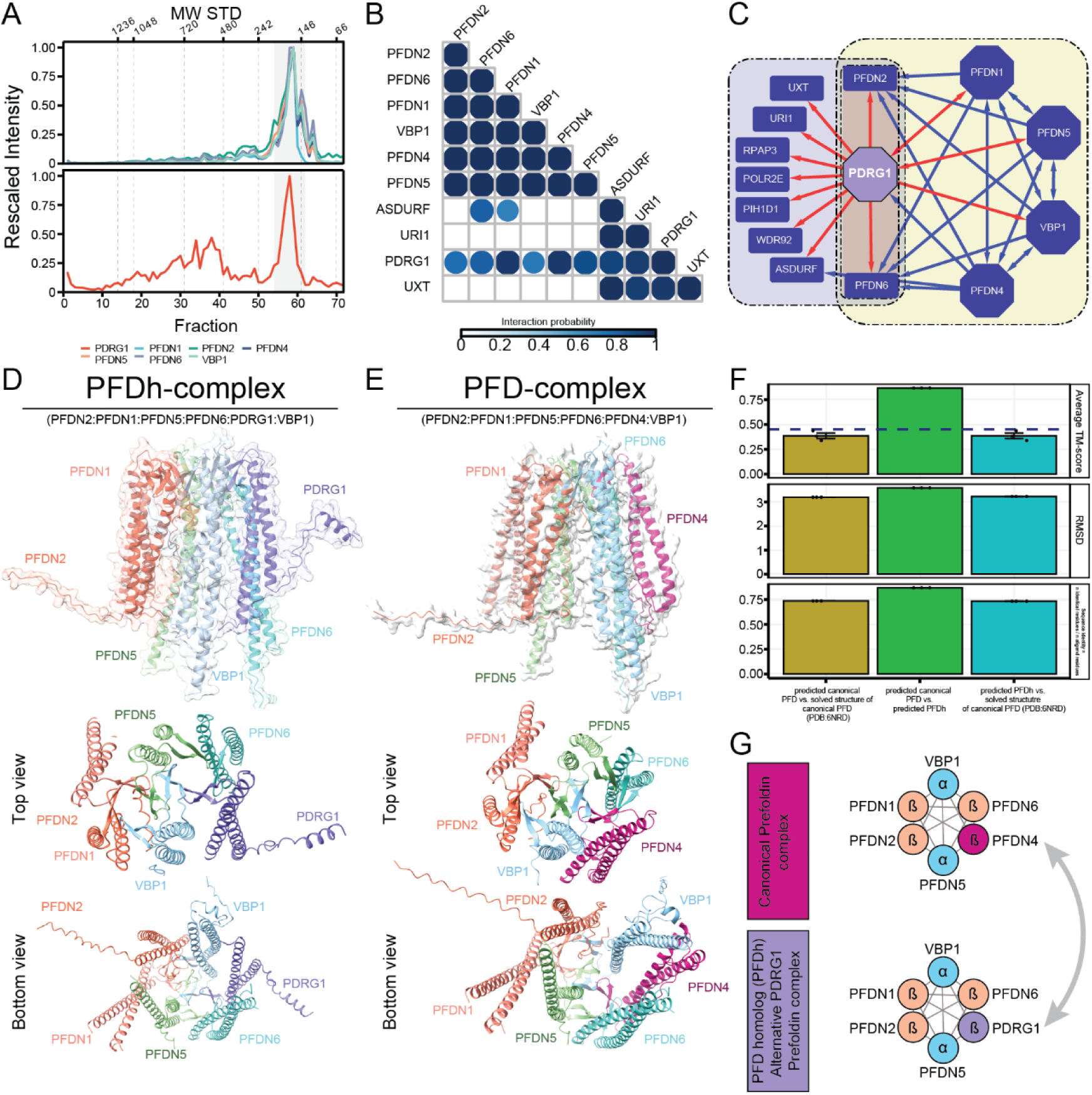
Data-driven identification of alternative assemblies and complexes in the PFD interaction network. **A**. PFDN2 DIP-MS coelution profile for canonical PFD complex (top panel) and PDRG1 (bottom panel). Y axis represents the normalized MS2 intensity across the replicates (n=3), while the X axis shows the fraction number. Molecular weight of the standards is highlighted on top of the plot. **B**. Triangular matrix representing the combined interaction probability derived from PFDN2 DIP-MS experiments (n=3) for the 10 core-subunits of the PFD and PFDL complex. **C.** PPI network from AP-MS data, using as bait proteins (octagons) mutually exclusive PFD subunits (PFDN1, VBP1, PFDN4, PFDN5) and PDRG1 (purple). PDRG1 interactions are shown in red. Canonical PFD complex (yellow background), and PAQosome subunits (blue) are indicated; Shared subunits (red grid background) are in the new proposed assemblies PFDN2 and PFDN6 expanded by PDRG1. **D.** Structural model of the PFD homolog complex containing PDRG1 instead of PFDN4 (complex subunits are colored). **E.** Structural model of the canonical PFD complex. **F.** Structural alignment of the two predicted models of canonical PFD and PFD homolog and an experimental PFD (PDB: 6NRD). The average TM-score indicates that the two predicted complexes show significant structural similarity (average TM-score > 45), and some structural similarity to the experimental structure. **G.** Proposed new PFD homolog complex containing PDRG1 instead of PFDN4.

**Fig. 5.**
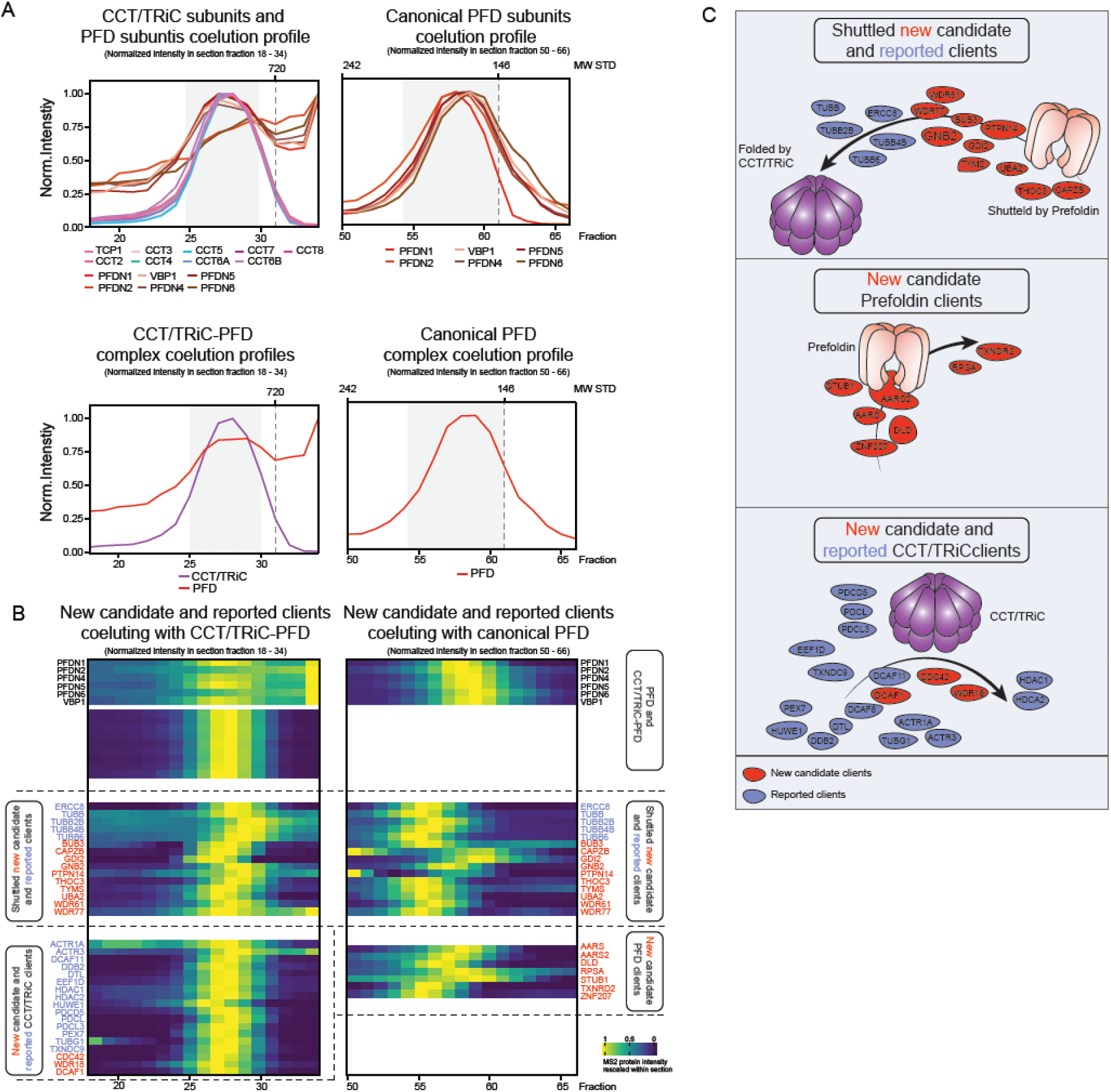
Assignment of new candidate clients for PFD and CCT/TRiC-PFD chaperone complexes. **A**. Coelution profiles of all CCT/TRiC-PFD subunits, the summed CCT/TRiC-PFD complex, the summed PFD complex, and the PFD complex subunits. Bottom X axis represents the fraction number, while the top X axis shows the molecular weight from the gel standards. Y axis shows the rescaled protein intensity **B.** The heatmap showcases reported (blue labels) and new candidate (red labels) clients are grouped in proteins coeluting with CCT/TRiC and PFD (potentially shuttled), PFD alone (folded by prefoldin) and CCT/TRiC alone (folded by PFD). Cell colors of the heatmap represents the MS2 protein intensity rescaled to the maximum intensity of each protein within the section of gel fractions represented. **C.** Graphical representation of the identified clients and their interacting complexes. Red circles represent newly identified candidate clients while blue circles indicate reported clients.

**Fig. 6.**
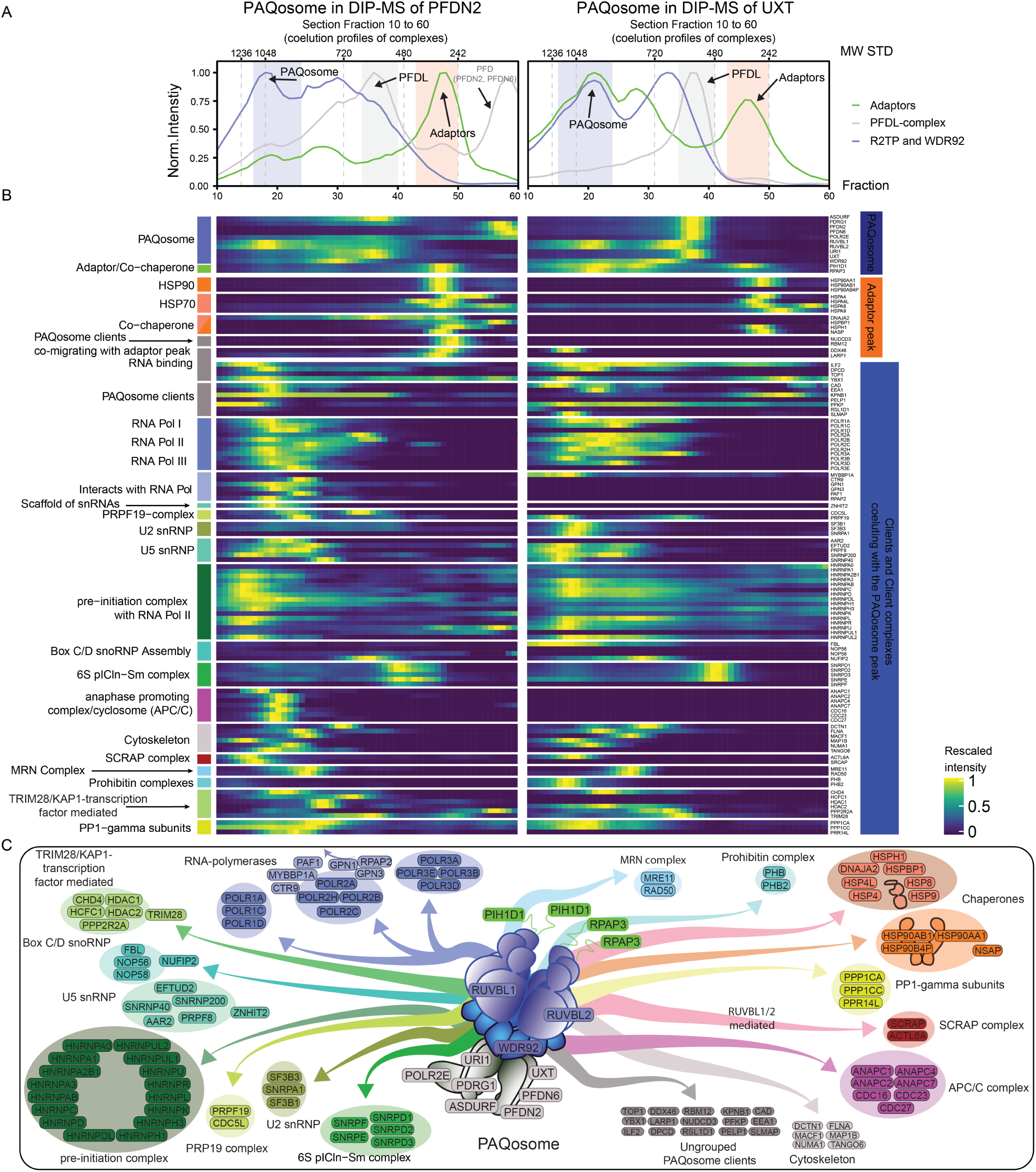
Assignment of PAQosome clients and client protein complexes. **A.** Coelution profiles for adaptors complex (green-line), PFDL-complex (grey-line) and R2TP module (blue line). X axis represents the fraction number (from 10 to 60) while the Y axis represents the rescaled intensity. Colored boxes represents either the PAQosome (blue box) or the adaptor peak (green box, RPAP3, PIH1D1). The top X axis shows the molecular weight from the gel standards. The two facets represent the two bait proteins employed. To investigate the peaks within the section, see Extended data Fig. 2 and the respective coelution group. **B.** Heatmap showcasing the identified clients within the UXT and PFDN2 DIP-MS data. The annotation on the right groups highlights if the protein is interacting with either to adaptor peak (orange) or the major PAQosome assembly (blue). The clients were further deconvoluted by literature-based knowledge into assemblies, which are indicated on the left side annotation. The color code corresponds with the graphical representation below. Cell color represents the rescaled intensity. **C.** Graphical representation of the identified PAQosome client complexes.

To understand in greater depth the organization of the PDRG1-containing prefoldin complex, which we dubbed PFD homolog (PFDh), we performed structural prediction of the PFDh and PFD complex using ColabFold [53]. Both heterohexameric complexes were predicted with high confidence, with a weighted confidence score of 0.8 for canonical PFD and 0.77 for PFDh (Supplementary Data S8), resulting in the ‘jellyfish’ structure formed by the stacking of the *α*2*β*4-prefoldin subunits into the typical *β*-barrels and the six protruding tentacle-shaped coils [54] (see Fig. 4D,E for predicted structures of the best ranking model of both complexes, see Extended data Fig. 5C-F for model confidence score and predicted aligned error (PAE)). Multiple structural alignments (MSTA) by US-align [55] of the two predicted structures and an experimental PFD structure showed a significant overlap of predicted PFD vs. the PFDh (TM-score: 0.869). Both predicted structures displayed weak similarity to the experimental determined PFD complex (PDB:6NRD) (Fig. 4F, Supplementary Data S8), due to the absent tails in the experimental structure. Of note, the PFDN4 to PDRG1 switch allows for the N-terminal tail to extrude from the predicted structure, potentially forming an additional *α*-helix in comparison to the canonical PFD. This feature could mediate new interactions by changing the substrates specificity of this complex. As a proxy for the abundance of the PFDh complex, the MS2 based intensity of PDRG1 in the PFD/PFDh peak was compared with the abundance all other PFD subunits (Extended data Fig. 5G and Supplementary Data S9). This shows that the PFD complex is 28.4 times more abundant than the PFDh complex, indicating that only 3.5 % of the peak can be attributed to PFDh complex (see Extended data Fig. 5H).

Thus, our data strongly suggests the presence of at least 2 similarly sized PFD complex isoforms not yet identified or reported in literature. More work will be needed to identify the molecular function of the newly discovered PFDh (Fig 4G).

### Identification of core PAQosome and PFDL components

The DIP-MS approach of PFDN2 and UXT also identified ASDURF as coeluting with PFDL and the PAQosome complex (Extended data Fig. 6A). In line with our data, this protein has been recently shown to be a component of the PAQosome [52]. To validate the presence of ASDURF in the PAQosome, we performed reciprocal AP-MS using the tagged versions of R2TP members (RUVBL1 and RUVBL2), the adaptors RPAP3 and PIH1D1 and the PFDL components UXT, PDRG1 and PFDN2 (Supplementary Data S7). The results indicated high-confidence interactions between ASDURF and all the tested bait proteins, thus validating the DIP-MS findings (see Extended data Fig. 6B). Binary structural alignments employing US-align [55] of the AlphaFold2 [56] predicted structures of ASDURF and the other prefoldin subunits showed a consistently high structural alignment TM-score for ASDURF versus β-prefoldin-subunits (see and Extended data Fig. 6C and Supplementary Data S10). In comparison to subunits of CCT/TRiC and R2TP-core and adaptor proteins, α- and β-subunits showed an average TM-score of 0.563 and an RMSD of 2.03 Å, indicating that ASDURF shares structural topology features with prefoldins. In contrast alignment to R2TP and CCT/TRiC subunits only had an average TM-score of 0.252 and an RMSD of 3.68 Å (Extended data Fig. 6D) indicating the absence of shared structural topologies. To obtain an acceptable structural model of the PFDL complex, we had to cut away the intrinsically disordered regions of URI1 (Extended data Fig. 6E), which allowed us then to model the PFDL complex, with a weighted confidence score of 0.75 (Extended data Fig. 6F and Supplementary Data S10, see Supplementary information for more details). Thus, our DIP-MS and AP-MS data, and orthogonal prediction of the structure of PFDL module, allowed us to conclude that ASDURF is included as PFDL subunit and to thus identify it as a component of the PAQosome supercomplex.

Lastly, the UXT DIP-MS data consistently identified a PPI between the PFDL complex and POLR2E which was often considered an associated protein for the PFDL-module or the PAQosome, but not a core-component. Of note, in all 6 DIP-MS experiments, regardless of the bait employed, we observed the PFDL complex interacting with POLR2E (coelution profiles in Extended data Fig. 6A). AP-MS analysis of any tagged PFDL like subunit (URI1, UXT, PDRG1, PFDN2) identified POLR2E, strongly suggesting that POLR2E is constitutively associated with the core subunit of the PFDL complex in-vivo. Accordingly, in previous work [28], it has been shown that URI contains a dedicated high-affinity POLR2E binding domain, suggesting that the PFDL complex is tightly linked to POLR2E in absence of other PAQosome members.

These results suggest that the *in vivo* stable form of the PFDL complex deviates from the hexameric paradigm derived from the PFD literature [57, 58]. We propose an updated configuration of the PFDL complex to a hetero-heptameric assembly, containing POLR2E.

### Identification of canonical PFD clients by DIP-MS

PFD together with HSP90, CCT/TRiC or as standalone complex has been reported to interact with several client proteins to mediate their folding [59]. These client-chaperone interactions are usually substoichiometric and not detectable by SEC-MS. We leveraged the increased sensitivity of DIP-MS towards low-abundant interactions to identify reported and putative novel clients of PFD and CCT/TRiC complex.

To gain insights into both complex-complex interactions (CCIs) and protein-complex interactions (PCIs), we utilized the PPI network derived from the *PPIprophet* toolkit (i.e. predicted interactions passing the 10% FDR threshold set up for every experiment) and selected the subnetworks corresponding to proteins interacting with CCT/TRiC and PFD (Fig. 5) or the PAQosome (Fig. 6). Within these subnetworks, we superimposed the available knowledge of the respective complexes to identify CCIs, while proteins absent from the complex databases were selected as PCIs.

We considered CCI’s as potentially high-order assemblies and specialized machines and PCI’s as putative clients. We further considered known structural features of distinct protein classes reported to be clients of CCT/TRiC (e.g., WD-repeats containing proteins) [60] (Supplementary Data S12).

Both canonical PFD and CCT/TRiC-PFD complexes showed excellent coelution of all subunits (Fig. 5A). Proteins and complexes such as actin, tubulin subunits and heat shock proteins multimers have been reported to be shuttled from PFD to CCT/TRiC for folding [18–20]. The DIP-MS results recapitulated these findings by detecting their coelution with both the PFD peak and the CCT/TRiC-PFD supercomplex peak (Fig. 5A, B). Thanks to the unbiased nature of our approach, we also identified novel putative clients. As example, the G-protein GNB2 has been previously identified as interaction partner of PFD subunits and PDRG1 [5, 61] adding orthogonal evidence to our previous discovery of the PFD homolog complex. We observed coelution of GNB2 with both the PFD complex and the CCT/TRiC-PFD, suggesting that GNB2 is most likely shuttled via PFD to CCT/TRiC. We validated this interaction by reciprocal AP-MS (Extended data Fig. 7A) and found additional evidence of the GNB2-PFD PCI in several large-scale AP-MS datasets such as BioPlex [5] (Extended data Fig. 7B) and in a study characterizing the human chaperone interactions with the canonical PFD subunits PFDN5 and PFDN2 [5, 61]. Interestingly, GNB2 and other G-proteins are folded by CCT/TRiC in an interplay with co-chaperone phosducin-like protein 1 (PDCL) [62, 63]. In the PFDN2 DIP-MS data we identified coelution for several phosducin-like proteins such as PDCL and PDCL3 with CCT/TRiC (Figure 5B, C). Overall, at the CCT/TRiC peak we identified 20 proteins showing coelution with CCT/TriC for which evidence of interaction with CCT/TRiC-subunits was reported [5, 64–66] and 3 which are not present in literature. It is important to point out that 15 of these are CCT/TRiC-exclusive interactions (i.e. showing coelution only with CCT/TRiC), and 5 are shuttled by PFDN to CCT/TRiC (i.e show coelution at both the CCT peak and the PFDN peak), which showcase the great degree of granularity achievable by DIP-MS (see summary in Supplementary Data S12). A comparison between the two DIP-MS experiments, shows that within the UXT DIP-MS experiment, only two thirds of the CCT/TRiC-PFD and PFD core-subunits were identified (Extended data Fig. 8A). CCT/TRiC is not an interactor of the UXT DIP-MS but, due to its abundance, has been repeatedly found as background in affinity enrichment-based experiments [67]. Thus, we expect that CCT/TRiC identified clients should be either absent in the UXT DIP-MS or less abundant compared to PFDN2 DIP-MS. For the clients, only 10 (17%) were quantified in the UXT DIP-MS experiment (Extended data Fig. 8B). The protein abundance of core-subunits of CCT/TRiC-PFD was lower at the coelution peak and within the DIP-MS of UXT not all prefoldin subunits were quantified (Extended Fig.8C). All the CCT/TRiC clients quantified in the UXT DIP-MS were recovered with lower abundance compared to the PFDN2 DIP-MS (Extended data Fig. 8D). Within PFDN2 the CCT/TRiC-PFD subunits are enriched on average by 7.95 log2FC (Extended data Fig. 8E), and the identified clients were enriched 6.04 log2FC compared to the UXT-DIP-MS (Extended data Fig. 8F and Supplementary Data S13). Overall, our data suggests that PFD specificity extends beyond the previously reported client complexes and suggest a broader PFD role in the stabilization of unfolded nascent proteins and their transport to CCT/TRiC for folding as reported in the model in Figure 5C. The presence of newly putative PFD complexoforms, such as the newly identified PFDh, could be partially responsible in broadening the spectrum of the Prefoldin clients.

### Large-scale identification of PAQosome client-complexes and clients by DIP-MS

Next, we queried our scored DIP-MS data from the UXT and PFDN2 bait proteins for client complexes and client proteins of the fully assembled co-chaperone complex PAQosome. The PAQosome is a large multiprotein assembly that assists the molecular chaperone HSP90 in the assembly of protein complexes, such as small nuclear ribonucleoproteins involved in m-RNA splicing (snRNPs U4 and U5) [68], the three RNA polymerase complexes [43, 69] and box C/D snoRNA assemblies [70]. Further, it was reported that the PAQosome stabilizes Phosphatidylinositol 3-kinase-related kinases (PIKKs) and many other client complexes [71]. Utilizing prior knowledge on the reported client complexes (75 proteins, 13 complexes, see Supplementary Materials and Methods and Supplementary Table S1), we identified 45 of these 75 clients in the UXT DIP-MS data, and 91% of which (n=41) were scored by *PPIprophet* as PAQosome interactors (Supplementary Data S14). In addition, we were able to assign to the PAQosome dozens of additional proteins (after *PPIprophet* prediction), i.e., putative candidate clients, not included in this list, as described below.

We observed that most of the target proteins coeluted with the major PAQosome peak (Figure 6A) or the adaptor peak formed by RPAP3:PIH1D1 (Figure 3B, Figure 6A). Based on the coelution and evidence from orthogonal AP-MS datasets (Supplementary DataS14), we assigned 92 proteins (organized in 19 complexes) to the PAQosome assembly and 15 co-chaperones (organized in 4 assemblies) to the adaptor peak (Figure 6B).

Among the known PAQosome client complexes, we recovered all three RNA polymerases (RNAPs), which were previously reported to interact with the PAQosome [43, 69]. Besides the core RNAP complexes, in the PAQosome peak we found additional polymerase associated proteins such as GPN-loop GTPase 1/3 (GPN1, GPN3) and RPAP2 which are reported to be assembly factors for RNA Pol II and associate with RNA Pol II prior to nuclear import [72, 73]. We also recovered multiple snRNA assemblies including U2, U5 and PRPF19 assembly, together with ZNHIT2 which was reported to regulate snRNA complex formation [31] (Figure 6C). Moreover, we identified the box C/D snoRNAs (U3) assembly, which was reported to be assisted in assembly by the PAQosome [70]. We additionally identified an assembly mediated by the transcriptional repressor KAP1/TRIM28, which was reported to interact with URI1. Thereby URI1 recruits a PP2A subunit which dephosphorylates TRIM28 leading to repression of TRIM28 regulated retrotransposons. And indeed, we identified a PP2A subunit (PP2R2A) and TRIM28, associated with the PAQosome [74]. Furthermore, we identified candidate client-complexes not yet reported for the PAQosome, such as the Prohibitin complex or the Anaphase-promoting complex (APC/C). For single APC/C subunits, studies previously identified several interactions for RPAP3 and PFDN2 [57, 75], but to our knowledge it was not reported that the PAQosome is associated with the entire complex. Although we did not identify the complex as significantly enriched within our AP-MS data, we recovered 8 PPIs between PAQosome subunits and APC/C subunits within our DIP-MS experiments, suggesting that this is a low-abundant interaction, only accessible through significant increase in analytical sensitivity.

In the lower MW coelution group (apparent MW of 356 kDA or fraction 45) consisting of the adaptor/regulatory subunits RPAP3 and PI1HD1 of the R2TP, we observed a coelution of HSP90-subunits (HSP90AA1, HSP90AB1 and HSP90B4P), which are reported to independently form a complex with RPAP3 at a ratio of 2:1 or 2:2 [76] (Figure 6B). Additionally, in the DIP-MS of PFDN2, HSP70 subunits (e.g., HSPBP1, HSPA1L) coelute in a similar MW range, showing additional client-chaperones for the two R2TP-adaptors [50, 76].

Finally, as the nature of our DIP-MS approach supports the recovery of prey-prey interactions, we recovered an additional 1117 PPIs between the client-complexes, exemplifying the high-density of the DIP-MS generated PPI network. For 22 grouped client-proteins, no direct PAQosome-interaction was previously reported in literature and for 4 of them (MACF1, PRR14L, RSL1D1, TOP1) we recovered PPIs within our reciprocal AP-MS dataset with PAQosome components such as RPAP3, PIH1D1 and UXT (Supplementary Data S7). It is important to point out that most of these peaks represent fractional amounts of the total protein signal as many proteins moonlight into different assemblies or show a CCI between a client-complex and the PAQosome.

Last we compared the abundance of the recovered PAQosome clients in the UXT and PFDN2 DIP-MS. The PAQosome core-subunits were fully identified in both DIP-MS experiments (compare Extended data Fig. 2). 102 clients (or 95%) were quantified in the PFDN2 DIP-MS versus 72 clients (or 68%) in UXT DIP-MS experiment (Extended data Fig. 9A, B). We found that 31 clients were enriched with at least 0.5 log2FC in the UXT DIP-MS and 54 clients in the PFDN2 DIP-MS experiment (log2FC UXT/PFDN2 DIP-MS) (Extended data Fig. 9C, Supplementary data S15). These indicates that the two DIP-MS experiments overlap across large-parts of the recovered clients, allowing to complement each other. The specificity of the above reported PAQosome clients has been demonstrated via the specific enrichment of PAQosome clients in the AP-MS using PAQosome subunits as baits compared to the enrichment observed with Prefoldin specific bait proteins (see Extended data Fig. 9D).

Overall, the identification of a large portion of reported clients for both the CCT/TRiC and the PAQosome as well as novel putative client complexes and client proteins demonstrates the resolution of DIP-MS for dissection of a particular interaction network of interest and the recovery of functionally relevant interactions in the context of cellular proteostasis.

## Discussion

The precise characterization of molecular networks and state-specific changes thereof has been a long-standing goal of molecular systems biology. Identification of core and accessory complex subunits, their variation in stoichiometry or composition across cell states and cell types and the interactions of these modules with substrate proteins or additional complexes (complex-complex interactions) is expected to significantly expand our knowledge on the proteome function and response in health and disease. In short, the context-specific modular organization of the proteome is of great biological significance and has remained analytically challenging.

Methods focusing on a single protein such as AP-MS, BioID, APEX or Y2H, while having high specificity, are fundamentally static and require numerous experiments to reliably identify protein complexes, which hinders their application across cell types, perturbations or time-points due to the exponential increase in resources needed.

Recently introduced co-fractionation methods such as SEC, IEX or WAX can quantify thousands of co-fractionating proteins and, potentially, deconvolute complexes within a single experiment. However, the dynamic range of protein expression and complexity of the interactome are such that co-fractionation approaches have so far not been able to obtain interaction networks at a resolution comparable to AP-MS or Y2H.

To overcome the limitations of AP-MS and co-fractionation MS, in this study we introduce a high throughput method dubbed DIP-MS to deconvolve the composition of affinity-enriched protein complexes, yielding insights into contextual protein complex organization at an unprecedented depth. We benchmarked DIP-MS versus the two state-of-the art MS-based techniques (AP-MS and SEC-MS) to resolve protein complex composition and showed that DIP-MS combines the specificity of AP-MS while benefiting from the larger number of (size-resolved) interactions recovered from fractionation-based approaches.

As DIP-MS experiments contain various level of information from PPIs and complex-complex interaction, to stoichiometry and subassemblies, we developed *PPIprophet* to facilitate the analysis of DIP-MS datasets. *PPIprophet* employs deep learning to predict PPIs in DIP-MS data and applies FDR correction using standard target-decoy competition to distinguish true interactors from spurious, coeluting proteins. As this approach uses coelution profiles derived from the size fractionation of affinity-purified complexes, identification of complexes is straightforward.

Other frameworks for analysis of SEC-MS data such as PCprophet and EPIC [77] may report different results. However, although these approaches rely to a lesser extent on previously reported complexes, they still employ prior knowledge for FDR control (PCprophet) or network pruning (EPIC) and are hence not suited for fully knowledge-free searches.

To demonstrate the applicability of DIP-MS to identify all types of interactions aforementioned (core, accessory, complex-client and complex-complex) we applied DIP-MS to the protein complex landscape of the Prefoldin (PFD), PFDL (Prefoldin-like) using PFDN2 and UXT as bait proteins. This system strikingly exemplifies the complexities of modular protein organization in the cell in that presents multiple complexes organized in high-order assemblies (i.e., complex-complex interaction) which have various preferred substrates and play a fundamental role in cellular proteostasis.

These two experiments recapitulated two and a half decades of previous prefoldin characterization and interaction studies covering more than 184 interaction studies in less than a week of MS measurements. The data recovered subassemblies like the R2TP complex, complex-complex interactions such as the CCT/TRiC-PFD or the R2TP/PFDL (PAQosome) and discovered an alternative Prefoldin complex containing PDRG1 as core-subunit (which we validated by AP-MS). The data further confirmed recently reported interactions such as PFDL-subunits with ASDURF and assigned POLR2E as an *in-vivo* constitutive subunit of the PFDL-complex.

Overall, we identified a large fraction of the previously reported PFD and PFDL client complexes and proteins and, ultimately, advanced our understanding on the modular organization of this section of the human interactome. Notably, prior PPI knowledge extracted from databases typically represents the cumulative contribution of different methods applied to diverse cellular models under disparate experimental conditions, hence constituting a generic, average network. On the other hand, DIP-MS allows to probe the state of a complex network with great depth in a specific cellular state. Accordingly, we observed a great degree of modularity in the PFDN2 and UXT derived network, highlighting the intricate nature of the human interactome and the need for methods capable of capturing it. We identified additional complex instances for which the specific functions and preferred client proteins yet need to be elucidated. The general feature of prefoldin complexes of two alpha-and four beta-subunits potentially allow for additional combinations, which would allow for a heterogeneous mix of different assemblies most likely having specialized functions or preferred substrates.

Even though DIP-MS outcompeted reciprocal AP-MS in terms of number of identified interactions, it is interesting to observe that the DIP-MS data did not recapitulate all the tested interactions. This can be due to the following reasons, i) signal dilution, following extensive biochemical fractionation may compromise accurate coelution profiling of low abundant proteins, ii) true sample/state specific differences or iii) the complex stability influenced by the separation conditions employed in the gel, which could result in loss of interacting proteins or disassembly of large supercomplexes into stable submodules.

Furthermore, while our scoring approach employs a decoy-based solution to the problem of coeluting proteins, development of more sophisticated statistical frameworks might be beneficial to further filter contaminants and increase specificity in complex identification (e.g., based on CRAPome [67]). With an increasing number of DIP-MS experiments analyzed and annotated, contaminant complexes could be more specifically separated from true interactors hence benefiting all interactome studies by providing assembly-state context for common contaminant proteins. We further have to stress that, while techniques achieving greater theoretical resolution such as cryo-slicing-BNP [78] have been developed, they require specialized equipment and great amount of know-how thereby being practically impossible to transfer between laboratories. On the other hand, the simplicity and accessibility of the DIP-MS workflow could be readily implemented and adapted in a variety of experimental setups.

DIP-MS allows the characterization of all bait-containing protein complexes from roughly ∼1 µg of purified bait-protein complex (quantified by heavy spike-in), which can be obtained from approximately 6 x 10^8 HEK293 cells ectopically expressing the bait protein (∼30 mg of protein lysate). For comparison, our in-house reciprocal AP-MS study of 11 PFDN/PFDL baits required five time more cells and did not cover several key complexes recapitulated by DIP-MS such as CCT/TRiC or the R2TP.

Since our high-throughput DIA-MS approach allows complex resolution for a single bait within roughly a day, this cuts acquisition time over previous co-fractionation studies by at least 3 fold [7, 79]. Further, the high-reproducibility of separation by native-PAGE will enable the probing of a selected bait protein complex landscape not only at steady state, but across different cellular states, which is difficult to achieve by techniques such reciprocal AP-MS. The bands in a native PAGE are highly reproducible and it is possible to load up to 9 experiments, or 3 conditions in triplicates, onto a typical commercial BN-PAGE.

We foresee the application of DIP-MS as a valuable approach for high resolution interactome studies. It will help guide structural studies by providing experimental evidence for the *in-vivo* existence of specific assemblies, which are convoluted within binary interaction data derived from Y2H or AP-MS. Finally, it will generate comprehensive interaction networks with minimal amounts of purified baits which could be further probed under perturbations thereby providing the opportunity to increase our understanding of the modular organization of the proteome and its dynamic organization into protein complexes. The increase in MS sensitivity is expected to significantly lower the input material require for DIP-MS experiments and unlock the possibility of clinical applications. It can be envisioned the derivation of cells representing the patient-specific disease and the application of endogenous tagging coupled to DIP-MS to probe interactome differences between healthy and disease tissue hence bringing us closer towards understanding the relationship between proteotype and phenotype.

## Author contributions

F.F. and M.G. conceived and designed the project. F.F. performed the experiments. M.H and F.U. provided critical input to experimental design and biochemical protocols. F.F., A.F., X.P. and W.F. acquired the mass spectrometry data. F.F. and A.F. conducted the data analysis and A.F. developed the software tool for data analysis. F.F. and A.F. generated the figures and wrote the original draft supervised by M.G., R.C. and R.A. All co-authors contributed in reviewing and editing the manuscript. M.G. and R.A. provided funding and resources to support the project.

## Code and predicted structure availability

*PPIprophet* is available freely for academic use at under MIT license on GitHub at https://github.com/anfoss/PPIprophet. The predicted structures, coelution data and *PPIprophet* parameters are deposited on Github (https://github.com/anfoss/DIP-MS_data).

## Acknowledgments

Figure 1 was created with BioRender.com. Ideas for graphical representation of Prefoldin, PFDL and major PAQosome components was taken from graphical representations of Marie-Soleil Gauthier, Philippe Cloutier and Benoit Coulombe. FF, FU and MG were supported by the Innovative Medicines Initiative project ULTRA-DD (FP07/2007-2013, grant no. 115766). MG has received support from the EU/EFPIA/OICR/McGill/KTH/Diamond Innovative Medicines Initiative 2 Joint Undertaking (EUbOPEN grant n° 875510).

## Supplementary information

Supplementary Figures S1–S7, captions for Supplementary Tables S1–S2 and Supplementary Material and Methods and references.

## Supplementary data/ data source for figures

- Supplementary Data S1: Absolute quantification, external calibration curve and inputs for PFDN2 and UXT.
- Supplementary Data S2: DIP-MS of PFDN2 and UXT (CCprofiler, min-max normalized, realigned, all three reps and summed gradient).
- Supplementary Data S3: List of assemblies and subassemblies of the PFD, PAQosome and CCT/TRiC subunits identified across the DIP-MS experiments.
- Supplementary Data S4: Abundance comparison of PFD, PFDL and PAQosome core-subunits across transcriptomics and co-fractionation approaches
- Supplementary Data S5: Identity matrix of sequence alignment of PDRG1 with PFDN4.
- Supplementary Data S6: Sequence alignment file of PDRG1 with PFDN4 in jvp-format.
- Supplementary Data S7: Reciprocal AP-MS of 11 core-subunits of the PFD and PAQosome complexes.
- Supplementary Data S8: Weighted scores for complex predictions of canonical PFD and PFD homolog. Structural alignment results.
- Supplementary Data S9: MS2 protein abundance comparison of PFD and PFD homolog to infer abundance of PFD homolog.
- Supplementary Data S10: Structural alignment of AlphaFold predicted structures of PFD and PFDL-subunits versus CCT/TRiC-subunits. Structural alignments of modelled PFDL complex to PFD and PFDh complex.
- Supplementary Data S11: URI1 sequence analysis to identify intrinsically disordered regions and protein binding sites.
- Supplementary Data S12: Result table of DIP-MS experiments with complex subunits and clients assignment for PFD and CCT/TRiC-PFD complexes.
- Supplementary Data S13: MS2 protein abundance comparison in coelution regions for PFD, CCT/TRiC-PFD and clients between the DIP-MS experiments of PFDN2 and UXT. The maximum signal at the coelution peak is used for comparison.
- Supplementary Data S14: Result table of DIP-MS experiments with complex, client-complexes and clients assignment for the PFDL containing PAQosome, PFDL and the Adaptor/HSP90-coelution groups.
- Supplementary Data S15: MS2 protein abundance comparison in coelution regions for the PFDL containing PAQosome, PFDL and the HSP90-adaptor
- groups between the DIP-MS experiments of PFDN2 and UXT. The maximum signal at the coelution peak is used for comparison.
- Supplementary Data S16: Molecular weight calibration curve realigned across all 6 DIP-MS experiments.
- Supplementary Data S17: Comparison of different filter plates for optimization of high-throughput DIP-MS sample preparation protocol.

## Material and Methods

More detailed information for material and methods are provide in the Supplementary information.

### Cell culture

Flp-In HEK293 T-REx cells ectopically expressing strep-HA tagged bait protein were essentially generated as described before by Glatter et al. [80]. Cells were cultured in DMEM (4.5 g/L glucose, 2 mM L-glutamine) supplemented with 10% FBS, 1% Pen-Strep, maintained at 37 °C and 5% CO_2_. HEK293 cells were grown on a 15 cm Nucleon dish (Thermo-Fisher) to reach 80% confluence before expression of the bait-protein was induced (1 *µ*g/mL doxycycline for 24 h). For harvesting, plates containing 2×10^7^ cells, were placed on an ice-cold rack and the media was removed. Cells were flushed off with 5 mL of PBS pH 7.4 (Gibco) and pelleted at 4 °C at 300×*g*. The supernatant was removed for subsequent shock freezing of the cells in liquid nitrogen and storage at −80 °C.

### Protein purification for AP-MS experiments

For 11 of 16 core-subunits of the PFD and PFDL containing PAQosome complexes, cell lines expressing SH-tagged-proteins were generated. These subunits included: the PFD/PFDL shared subunit PFDN2; canonical PFD subunits PFDN1, PFDN2, VBP1, PFDN4 and PFDN5; and the R2TP complex subunits RUVBL1, RUVBL2, RPAP3 and PIH1D1; and UXT, a PFDL/PAQosome subunit not shared with the other complexes.

For each AP a cell pellet derived from four 150 mm culture dishes (Nucleon Delta, ThermoFisher Scientific) was lysed in 4 mL ice-cooled HNN-lysis buffer (50 mM HEPES, 100 mM NaCl, 50 mM NaF, pH 7.4) supplemented with protease inhibitor cocktail (Sigma), 1 mM PMSF, 400 nM Vanadate, 1.2 *µ*M Avidin and 0.5% NP40 (Ipegal, Sigma). Lysed cells were rested on ice for 10 minutes before clarification by centrifugation at 16’000×*g* at 4 °C for 20 minutes. Strep-Tactin Sepharose (IBA) beads were equilibrated with 2 column volumes of HNN-lysis buffer and resuspended at 50% (v/v) slurry in HNN lysis buffer. The lysates were incubated with 100 *µ*L of beads slurry on an end-over-end tube rotator at 12 rpm for 45 minutes. Cleared lysate was loaded on Bio-Spin chromatography columns (Bio-Rad), to separate supernatant from the beads. Beads were washed twice with 1 mL ice-cooled HNN-lysis buffer and thrice with 1 mL of HNN buffer without supplements. Purified complexes were eluted by addition of three times 200 µL 2 mM Biotin (IBA) buffer. To precipitate enriched proteins, trichloroacetic acid (TCA) was added to 25% (v/v) and samples were incubated overnight at -20 °C. The proteins were pelleted by centrifugation at 16’000*×g* for 20 minutes at 4 °C and the supernatant was removed. The pellet was washed twice with 900 *µ*L -20 °C cold Acetone (Thermo Scientific) with centrifugation of 16’000*×g* at 4 °C for 5 minutes to remove remaining TCA. After removal of the supernatant, pellets were dried under vacuum for 10 minutes.

### MS-sample processing of reciprocal AP

The dried protein pellet was resuspended in in 30 *µ*L of 8 M Urea dissolved in 100 mM Ammonium bicarbonate (ABC) (Fluka) pH 8.8 and incubated at 1000 rpm at 25 °C on a thermoshaker (Eppendorf). Urea concentration was lowered to 0.8 M by addition of 250 *µ*L of 100 mM ABC. For reduction of disulfide bonds, TCEP (Aldrich) was added to a final concentration of 5 mM and the samples were incubated for 30 minutes at 37 °C and 1000 rpm. Ioadacetamide (IAA) (Sigma) was added to 10 mM final concentration for alkylation of cysteine residues and the samples were incubated in the dark for 45 minutes at 25 °C at 1000 rpm. Proteins were digested by addition of 1 *µ*g of Trypsin (Promega) and incubated overnight at 37 °C at 600 rpm on a thermoshaker. Proteolysis was quenched by addition of 0.05% TFA (Thermo Scientific) to reach ≈ pH 2. Peptides were purified over a C18 microspin column (Nest Group). Briefly, C18 columns were activated by washing twice with 200 *µ*L 100% MeOH (Fisher Scientific) and twice with 200 *µ*L 80% ACN (Fisher Scientific) in 0.1% TFA (Fisher Chemical). Columns were equilibrated trice with 200 *µ*L 2% ACN, 0.1% TFA in H_2_O, before loading the samples. The bound peptides were washed 5 times with 200 *µ*L 2% ACN, 0.1% TFA in H_2_O before elution with a 100 *µ*L of 50% ACN in 0.1% TFA. Peptides were vacuum dried and stored at -80 °C.

### Purification of complexes for Blue native-PAGE separation (BNP)

The affinity purification of protein complexes for native size-based fractionation followed the protocol described above for AP-MS samples, deviating in the following points. For a DIP-MS sample, thirty confluent 150 mm plates were used for one replicate. After lysis in HNN-lysis, the samples were treated with 5’625 Units of Benzonase (Sigma-Aldrich) and incubated at 10 °C at 500 rpm for 45 minutes. The cell lysate was clarified at 16’000×*g* at 4 °C for 20 minutes. 30 mL of cleared lysate were separated into 7.5 mL aliquots and transferred to four 15 mL falcon tubes. For each tube 200 *µ*L of equilibrated 50% Strep-Tactin sepharose bead slurry was added and subsequently incubated on an end-over-end rotator at 12 rpm for 45 minutes. The resulting elution volume of 600 *µ*L per replicate were pooled together and concentrated over a 30 kDa MWCO filter (Amicon) at 4 °C and 3000×*g* to 35 - 50 *µ*L.

### Separation of co-purified complexes by blue native-PAGE

To separate co-purified protein complexes, 35 - 50 *µ*L of the concentrated sample were loaded on a blue native-PAGE (BNP). The native separation procedure followed previous protocols [13] for native enriched protein complex separation. First, the concentrated eluate was mixed with a 1:4 ratio with native gel loading buffer by carefully pipetting slowly up-and down. From this mixture 45 *µ*L were loaded with gel loader tips on the NativePAGE 3-12% Bis-Tris precast protein gels (Invitrogen). For replicate one and two of PFDN2 aliquots of 1 *µ*L and replicate 1 of UXT an aliquot of 0.5 *µ*L was taken away for absolute bait protein quantification. As molecular weight standard 10 *µ*L of NativeMARK MW-standard (Invitrogen) was added in the first lane. As cathode buffer a Light Blue Cathode Buffer was applied, else the manufacturer’s instructions were followed. The BNP was run for 3 hours at 4 °C with constant voltage: 120 V for 25 minutes, 160 V for 2h and 5 mins and 200 V for 30 minutes.

### MS-sample preparation of gel slices

The native-PAGE gel was rinsed with dionized H_2_O (diH_2_O) before washing it three times with 30 mL H_2_O for 5 minutes in a gel-box. Following the initial gel washing step, the gel was stained for 1 h with SimplyBlue^TM^ SafeStain. After removal of the staining solution, the BNP was rinsed with diH_2_O before destaining overnight in diH_2_O. Gels were imaged with a Fusion FX (VILBER), before slicing. The MW standard was noted on a millimeter paper and the line of each replicate was vertically cut to separate each replicate from the other replicates. An in-house designed gel-slicing tool with hundred 1 mm distanced racer blades mounted on a metal frame, was applied to each lane.

The slices were transferred from the gel-slicing tool to a 96-well glassfiber (GF) filter plate (Pall), which contained 200 *µ*L H_2_O. The filter plate was equilibrated by washing twice with 200 *µ*L of 100% acetonitrile (ACN) followed by one wash of 200 *µ*L of 50% methanol (MeOH) in 20 mM ammonium bicarbonate (ABC). The washing solutions were removed by centrifugation at 700×*g* for 5 minutes at room temperature. The position of the gel slices on the filter plate were randomized, to avoid processing biases. The gel slices were first de-stained by addition of three times 200 *µ*L 50% MeOH in 20 mM ABC followed by two washes with 200 *µ*L of 100% ACN to the gel slices with 5 minutes of incubation before each centrifugation step. Reduction was performed by addition of 50 *µ*L of 25 mM TCEP in 20 mM ABC at 90 rpm at 37°C for 30 minutes. Alkylation was conducted by addition of 50 *µ*L of 50 mM IAA in 20 mM of ABC and incubation in the dark at room temperature for 45 minutes. The slices were washed once with 200 *µ*L of 50% ACN in H_2_O, and then twice with 200 *µ*L of 100% ACN. To each well 50 *µ*L of the digestion mix, containing 0.5 *µ*g Trypsin (Promega), 0.1 *µ*g Lysyl Endopeptidase (FUJIFILM Wako) and 0.01% ProteaseMax^TM^ (Promega) in 20 mM ABC was added. After 25 minutes of incubation at 37 °C at 100 rpm, an additional 100 *µ*L of 20 mM ABC was added, to cover all gel-slices. Protein digestion was performed overnight at 37 °C with 100 rpm. In order to avoid evaporation of the digestion mix, the filter plate was closed with parafilm at the bottom and on top with a metal cover lid. Peptides were collected by centrifugation at 700×*g* for 5 minutes and transferred to LoBind tubes (Eppendorf). The filter plate was washed once with 100 *µ*L 50% ACN in H_2_O followed by a wash with 100 *µ*L of 100% ACN. The washing solutions were pooled with the collected peptides. Samples were dried at 45 °C on a vacuum drier and stored at -80 °C until MS-acquisition.

### Data acquisition of reciprocal AP samples

The reciprocal AP-MS samples were acquired in DDA-mode on a Q Exactive^TM^ Plus Hybrid Quadrupole-Orbitrap^TM^ Mass Spectrometer (ThermoFisher Scientific) interfaced with an Easy NanoLC 1000 HPLC equipped with an autosampler (ThermoFisher Scientific). The dried peptides were dissolved in 20 *µ*L of buffer A (2% ACN and 0.1% FA in H_2_O) with 1:50 (v/v) iRT peptides (Biognoysis). For each sample, 2 *µ*L were injected, and the peptides were separated by reverse-phase chromatography on a high-pressure liquid chromatography (HPLC) column (75 *µ*m inner diameter, New Objective) manually packed with 15 cm of C18 beads (ReproSil-Pur 120 Å, C18-AQ 1.9 *µ*m, Dr. Maisch GmbH) with a 10 *µ*m fused silica tip emitter. Peptides were separated over a 90 min gradient from 5% to 30% buffer B (95% ACN in 0.1% FA in H_2_O) in Buffer A followed by step increase from 30% to 90% of buffer B in 7 minutes and a column clean-up by holding the gradient for 8 minutes at 90% buffer B. The capillary voltage was set to 2.2 KeV and 250 °C. The MS was operated in positive mode, with MS1 and MS2 scans collected in the Orbitrap analyzer (OT/OT). MS1 resolution was fixed at 70’000 at 200 *m/z* while the resolution for fragment ions was set at 35’000. The MS1 scan range was set from 350 to 1650 *m/z* with a parent ion isolation window of 1.5 *m/z*. The default charge state was set to +2, and unassigned charge states and +1 were excluded from fragmentation. The AGC target for MS1 scans were set to 1e^6^ with a maximum accumulation time of 54 ms. The 15 most abundant precursors per scan were fragmented with high-collisionally induced dissociation (HCD) using a normalized energy of 27% and were subsequently excluded for 15 seconds. Fragments were accumulated in the Orbitrap until reaching 5e^4^ charges for a maximum of 120 ms.

### DIA data acquisition of native-PAGE separated AP samples

The DIP-MS samples were acquired in DIA-mode with a Q Exactive^TM^ Plus Hybrid Quadrupole-Orbitrap^TM^ Mass Spectrometer (Thermo) interfaced with the EvosepOne system. First, the dried peptides were dissolved in 250 *µ*L of buffer A (0.1% formic acid in H_2_O) (Thermo Fisher Scientific) with 1:2500 (v/v) iRT peptides (Biognoysis). Dried peptides were sonicated for 10 minutes, and centrifuged at 16’000×*g* for 10 minutes. To avoid loading small gel pieces, which might pass the glassfiber filter, 230 *µ*L of the 250 *µ*L were loaded on equilibrated Evotips. The C18 material of the Evotips was activated with 10 *µ*L of Buffer B (98% acetonitrile and 0.1% formic acid in H_2_O) and by soaking the tips in Propan-2-ol. Next, the tips were equilibrated by adding 10 *µ*L of buffer A, following by the addition of 230 *µ*L of resuspended peptides per fraction. Loading was completed by a centrifugation at 300×*g* for 5 minutes. To prevent drying of the C18 material, 200 *µ*L of Buffer A were added on top of the tips.

Peptides were separated on a fused silica PicoTip^TM^ with an inner diameter of 100 *µ*m (New Objective, Woburn, USA) and 50 *µ*M tip diameter, in-house packed with 8 cm of C18 beads (MAGIC, 3 *µ*m, 200 °A, Michrom BioResources, Auburn, USA). The peptides were separated using the ’60 samples per day’ method (24 minutes gradient for PFDN2 and 21 minutes gradient for UXT as bait protein) using the EvosepOne system. For data acquisition, the mass spectrometer was operated in positive mode with the capillary heated at 275 °C and maintained at 2.5 KeV. We employed for DIA a 22 variable windows with +1 DA overlapping on the upper window boarder, ranging from 350 to 1650 *m/z*. The full MS1 scan was performed over a mass to charge range of 150 to 2000 *m/z* with a high-resolution of 70’000 fixed at 200 *m/z*. The AGC target was set to 3e^6^ with a maximum accumulation time set to 200 ms.

For MS2 scans the resolution was fixed to 17’500 with an AGC target of 2e^5^ with HCD fragmentation in stepped mode employing a collisional energy of 25%, 27%, 30%, normalized to 500 *m/z* at charge state +1. Each MS2 scan was set to 50 ms, leading to a total cycle time of 1.3 sec.

### Reciprocal AP-MS data processing

The reciprocal AP-MS samples were analyzed with MaxQuant (version 1.5.2.8) and the built-in search engine Andromeda [81]. Raw files were searched against a human protein database downloaded from UniProtKB [82] (downloaded on the 01.12.2019) and supplemented with the protein sequence of GFP and the sequence of the SH-quant peptide sequence (AADITSLYK) [51]. For the search and subsequent quantification, the MaxQuant contaminant list [81] was included. The peptide identification search was performed with default parameters. Briefly, carbamidomethylation on cysteine residues was selected as fixed modification while oxidation on methionine residues and acetylation on N-terminal end were used as variable modifications. The maximal number of modifications were limited to five. The protein sequence database was searched for trypsin specific peptides, allowing up to 2 missed cleavages. For label-free quantification, the LFQ tab was enabled, applying the default parameters. Further, re-quantification and match between runs, with the default settings, were enabled. The peptide and protein false discoveries were controlled by 1% FDR respectively. For this purpose, decoys were generated using reversed sequence.

### Post-analysis of AP-MS data

The MaxQuant ’proteinGroups’ table was filtered (removed contaminants list identifiers from MaxQuant contaminant file) before Saint Express scoring. First reversed sequences, which passed the FDR were removed from the results (29 identifiers). The additional GFP control originating from a high pH fractionated GFP sample was added (see Supplementary Information). The final matrix was uploaded to CRAPome [67] to perform SAINT Express [83] scoring on spectral count level. Each of the 11 baits was scored independently against the GFP controls, as their interactomes are largely overlapping and scoring them together reduces the number of interactions recovered. For SAINT Express default parameters were used, with the adaptions that 10 virtual controls for the fold change calculation were applied and 4000 iterations and normalization for SAINT scores calculation. Next, the scored interactions were filtered by applying a Log2FC (FC_A_) ≥ 2 with a Saint probability ≥ 0.95. Further only interactors with an average spectral count ≥ 5 were kept, which removed low-abundant and inconsistently detected preys. Secondly, the filtered interaction partners were filtered against the most frequently detected background proteins across the CRAPome dataset, applying a 30% frequency threshold (excluding from the CRAPome some well-characterized PFD/PFDL interactions CCT/TRiC-subunits, HSP90, and TUBB2B, as these preys are with high-frequency and abundance detected in negative controls). This resulted in 407 binary interactions from 174 interaction partners across the 11 baits, which we categorized into 278 high-confidence interactions (HCI) from 99 preys (Log2FC ≥ 5 and Saint score ≥ 0.99) and 140 medium-confidence interactions (MCI) from 94 preys (Log2FC ≥ 2 and Saint between 0.95 and 0.99). Note the total sum of preys is higher than 174; depending on the bait, some preys can be alternatively classified as MCI or HCI. The resulting PPI-network is represented in Supplementary Fig. S5.

### DIP-MS data analysis

The DIA data were searched with Spectronaut (version 13.12.200217.43655, Laika), employing library-free directDIA against the human protein FASTA database downloaded from UniProtKB [82] (downloaded on the 01.12.2019) and supplemented with the protein sequence of iRTs [84] and the sequence of the SH-quant peptide [51]. The fasta file contained in total 20’366 entries, which were supplemented by decoy sequences within the spectronaut workflow. The analysis was conducted with default (BGS Factory settings) parameters with minor adaptions. Briefly, the peptide identification search was performed for tryptic peptides, allowing for up to 2 miss cleavages and a maximum length of 52 and a minimum length of 7 aminoacids. Carbamidomethylation of Cysteine residues was set as fixed modification (+57 Da) and N-terminal acetylation and oxidation on Methionine residues as variable modifications. A maximum of 5 modifications per peptide was allowed. Precursor and protein Q-value cutoff was set to 5%. For quantification the cross-run normalization and the best N fragments per peptide parameters were disabled. Quantification was performed on MS2 level, and the mean peptide quantity from all quantified fragments per stripped peptide sequence was reported.

### Post-processing of DIP-MS data

From the Spectronaut analysis, protein accessions, stripped peptide sequences, and peptide quantities per each fraction for each replicate were exported. Each DIP-MS replicate was processed in R using the filtering functions of the CCprofiler [16] R-package. We first filtered within each gradient the noisy peptide profiles by applying a consecutive protein ID based stretch filtering of 2 fractions for each peptide trace, which removed inconsistent quantified peptides. In addition, all non-proteotypic peptides were removed. Next, sibling peptide correlation was performed, to remove all peptides which do not show coelution across the separation range on peptide level. An absolute sibling peptide correlation cutoff of 0.2 was applied. After signal processing on peptide level, protein quantities were inferred by using the top2 highest intense peptides per protein. The protein matrices was used for visualization of protein complexes and serve as input for the *PPIprophet*. This conservative filtering and quantification parameters ensured (i) no noisy single hit wonders are used for PPI-identification or complex mapping, and (ii) that the intensities for each protein are comparable against each other.

### *PPIprophet* implementation

#### Quantitative protein matrices pre-processing and feature engineering

Training set was build using 33 datasets encompassing different separation techniques, number of fractions and organisms for a total of 1’675’356 PPIs [37]. Multiple organisms and separation techniques were employed to maximize the model generalization capabilities. Positive PPIs were derived from STRING (STRING combined score > 600) [85] while, to obtain the negative labels, random protein pairs showing weak correlation were used (correlation between -0.3 and 0.3) leading to a balanced dataset between positive and negative interactions. Protein profiles were smoothed using 1D discrete Fourier transformation and missing values were filled with the average value between the two-neighboring fraction. Following data smoothing and missing value imputation, the intensity vector was rescaled in a 0-1 range. To have fixed size input for learning we employed linear interpolation to rescale the fraction number to 72. For training 2 types of continuous features were calculated, similarly to the ones employed in our recently introduced PCprophet toolkit [86]. The features employed by *PPIprophet* are: i) sliding-windows correlation (w=6 fractions) ii) fraction-wide difference between protein intensity resulting in 2*n* features and 144 features when *n* = 72.

#### Deep learning model construction and training

Following data annotation, a deep neural network was constructed in Python v3.8 in Keras (https://keras.io) using Tensorflow2 (https://www.tensorflow.org) as backend. Input layer size was fixed to the number of features (144). For all other 3 layers, 72 neurons were used with Rectified Linear Unit (RELU) as activation function. To avoid overfitting 30% dropout was used for the hidden layers. In the final layer, sigmoid activation was used to classify coeluting and not coeluting PPIs. The model was trained using ADAM (learning rate = 0.001) and binary cross-entropy as loss function. To further mitigate overfitting, label smoothing of 0.1 was applied. The dataset was split in training and testing set using a 80:20 split and, the training set was further split in training and validation set using 70% of the data for training and 30% for validation. Model was trained for 256 epochs using a batch size of 64. EarlyStopping (patience=10) was used to avoid learning plateau and the best model was selected based on lowest validation loss, which was calculated after every epoch. Achieved performance metrics on the test set are reported in Table 2.

#### PPIprophet analysis

For new data a correlation matrix between all protein pairs is computed and non-negatively correlating pairs are then used for feature construction and deep learning prediction. For every protein a decoy PPI is generated by random selection of protein pairs absent from the target set previously generated. After generation of both target and decoy PPIs, features are calculated as previously described and the DNN model is used to discriminate coeluting and not coeluting PPIs. Following prediction, PPI probabilities from the DNN model are converted to empirical p-values and false discovery rate is controlled using the following formula [87].

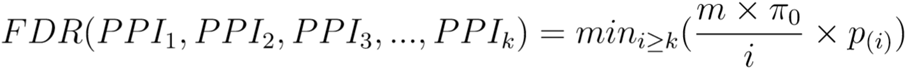

Where *π*_0_ is the probability that a putative discovery is false, *k* is the total number of selected discoveries and *m* is the number of putative discoveries, where for discovery is intended a PPI above the current probability interaction threshold. For every experiment an FDR cutoff of 10% was used. If replicates are present, the prediction probabilities for a particular PPI are combined into a weighted joint probability across replicates under the assumption of independence between the different replicates.

Following joint probabilities calculation, a combined adjacency matrix is generated where every edge is represented as the joint probability for that specific PPI. This combined adjacency matrix can be thought of as a series of *in silico* purification experiments where every column is a bait and every row is a prey

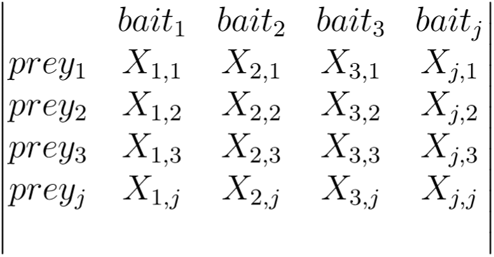

The score W for the interaction *bait*_1_*prey*_1_ is calculated assuming independence of prey and bait interaction from other interactions and is performed in vectorial format for computational efficiency.

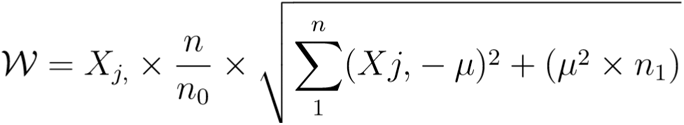

Where *X_j,_* represents the *jth* column, *n* is the total number of elements in the *jth* column, *n*_0_ and *n*_1_ respectively represents the number of negatively predicted interaction (probability *<* 0.5) and positive predicted interaction (probability > 0.5 and FDR lower than the user-set target FDR) in the *jth* column. The variable *µ* represents the average probability in the *jth* column and is defined as:

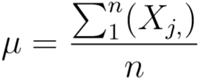

thereby *µ* intrinsically represents the specificity of the bait *j*. The term 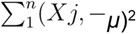 represents the square error compared to the bait which translates into penalizing proteins having similar probability to the average in the column, with the rational that true interactions have low *µ* and high square error *µ*. Following conversion of combined probabilities to scores a bootstrap procedure is applied to threshold the scores and further filter the interactions.

### Benchmark versus reciprocal AP-MS and SEC-MS

Each dataset (reciprocal AP-MS, PFDN2 DIP-MS and SEC-MS) was analyzed separately using SAINTexpress, *PPIprophet* or CCprofiler respectively using different thresholds for each of the tools. In this regard, we employed a strict 0.99 threshold for SAINTexpress (correspondent to a 0.01 FDR) as outlined in Post-analysis of AP-MS data, a threshold of 10% FDR for *PPIprophet* and a CCprofiler qvalue of less than 1% as reported in our previous study [16]. The CCprofiler derived complexes were converted to interaction network and used as is while for *PPIprophet*, positively predicted PPIs were selected and used directly. For comparison of abundances between SEC-MS and DIP-MS, for all proteins from the target list identified in the two experiments we selected the most abundant peak and averaged across replicates if present.

### Data analysis for DIP-MS of PFDN2 and UXT

To calculate ratios between Prefoldin, Prefoldin-like and the PAQosome complex, the protein elution profile of PFDN2 respectively UXT was used. Following peak selection, we assigned manually the PFD and PFDL peak and calculated the full width half max. The peak area was integrated using the trapezoid rule and divided by the entire PFDN2 or UXT signal across the entire fractionation dimension. For all stoichiometry calculations, subunits in each complex were selected and the full width half max was calculated. Then, the protein having the lowest area was used as stoichiometric unit. Each replicate was processed individually, and the bar-plot shows the data of 6 replicates from 2 independent DIP-MS experiment (3 PFDN2 and 3 UXT).

### Sequence alignment and prediction of intrinsically disordered regions

Sequence alignment of PFDN4 and PDRG1 was performed on the canonical FASTA sequence obtained from UniProtKB (3/7/2022) with Clustal Omega (EBI, version: 2.1)[88] using default parameters. The obtained sequence identity matrix is in Supplementary data S5. For visualization Jalview (version 2.11.2.0) [89] was used (sequence alignment file in Supplementary data S6). For the prediction of intrinsic disordered regions of URI1, the webserver of flDPnn (putative function- and linker based Disorder Prediction using deep neural network) [90] under http://biomine.cs.vcu.edu/servers/flDPnn/ was used, applying default parameters (26.06.2022). Outputs were limited to the relevant data containing predictions for disordered regions and protein binding interface.

### Protein complex structural predications

Structural prediction of alternative PDRG1 containing Prefoldin complex (PFD homolog, PFDh) were performed in ColabFold (version 1.3.0) [53], which combines MMseqs2 based multiple sequence alignment (MSAs) with AlphaFold2 [56] and AlphaFold-multimer [91] for protein complex structure prediction. Within ColabFold the MSAs were generated by MMseqs2 using the UniRef100 and environmental sequences. As input for the multiple sequence alignment the consensus sequences from the UniProtKB database were provided. As a template for the subunit order the deposited experimental structures of canonical PFD (PDB: 6NRD) [29] was used. The subunit order for PFDh complex and canonical PFD complex were the following: PFDN2:PFDN1:PFDN5:PFDN6:PFDN4/PDRG1:VBP1; and for PFDL: PFDN2:ASDURF:UXT:PFDN6:PDRG1:URI1(residue 24 to 222). For URI1 the disordered regions from residue 1 to 23 and residue 223 to 431 were removed from the URI1 prediction. For the multiple sequence alignment, the option for unpaired+paired setting was employed. Template mode was set to PDB 70. Under the advanced settings the default parameters were selected. The PFD and PFD homolog complex structure were relaxed (amber checked), but for PFDL the setting was not chosen due to time-outs during the prediction. Five models were generated per complex, whereas they were ranked by the weighted score. Weighted scores were calculated by the addition of 0.8 x pTM with 0.2 x ipTM-score (see Supplementary data S10 and S12). Structures and structural alignments were visualized with ChimeraX (version 1.4) [92]. All predicted complex structures are deposited in GitHub at https://github.com/anfoss/PPIprophet.

### Structural alignment of predicted structures

For binary structural alignments of ASDURF to other core-subunits, the tool US-align [55] (Version 20220511) was used (https://zhanggroup.org/US-align/). AlphaFold predicted structures were obtained from UniProtKB. The alignment was performed applying sequence independent structure alignment on the CA (protein) backbone of residues. TM-scores, normalized to the length of ASDURF, and the second protein, RMSD and alignment length (Supplementary data S12). A threshold of > 0.5 on the average TM-score was used to estimate if the structure of ASDRUF shares same global topology with other core-subunits of the PFD, PFDL and PAQosome subunits. CCT/TRiC subunits alignment were used as negative controls. Alignments of predicted complex structures (canonical PFD, PFD homolog and PFDL complex) were performed by multiple structure alignment (MSTA) using US-align with default parameters and a TM-cutoff of 0.45. As control an experimental canonical PFD structure (PDB: 6NRD) was used, although it lacked sufficient resolution.

## Data availability

The mass spectrometry proteomics data have been deposited to the ProteomeXchange Consortium via the PRIDE partner repository with the dataset identifier PXD035032.

## Key Resources Table

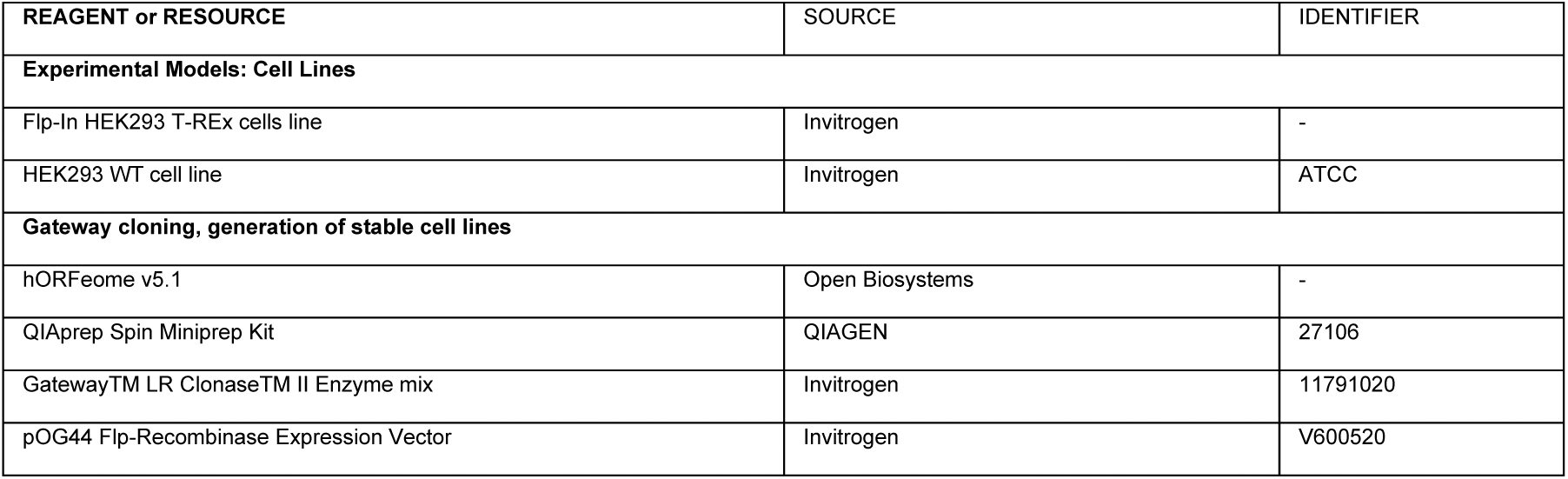
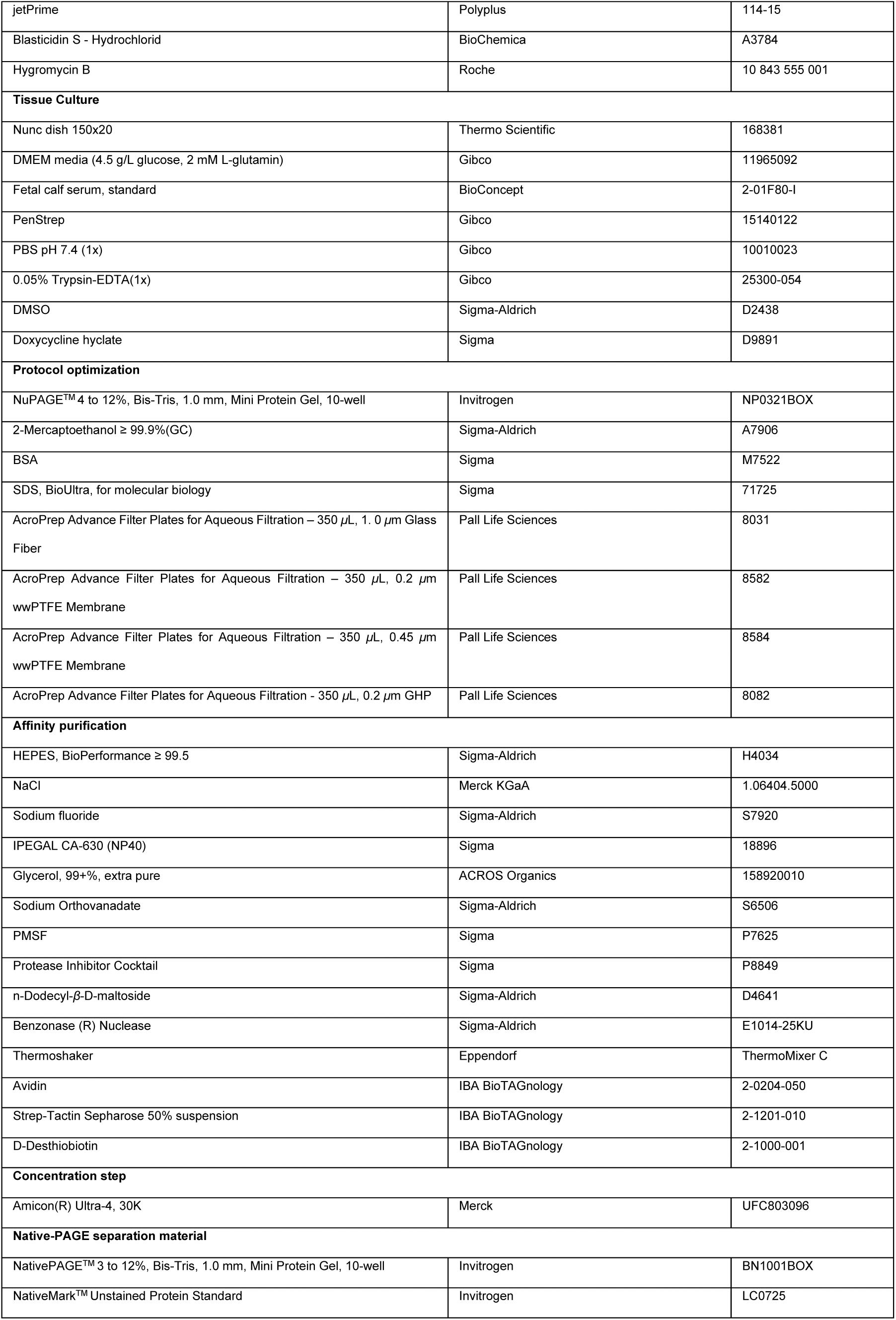
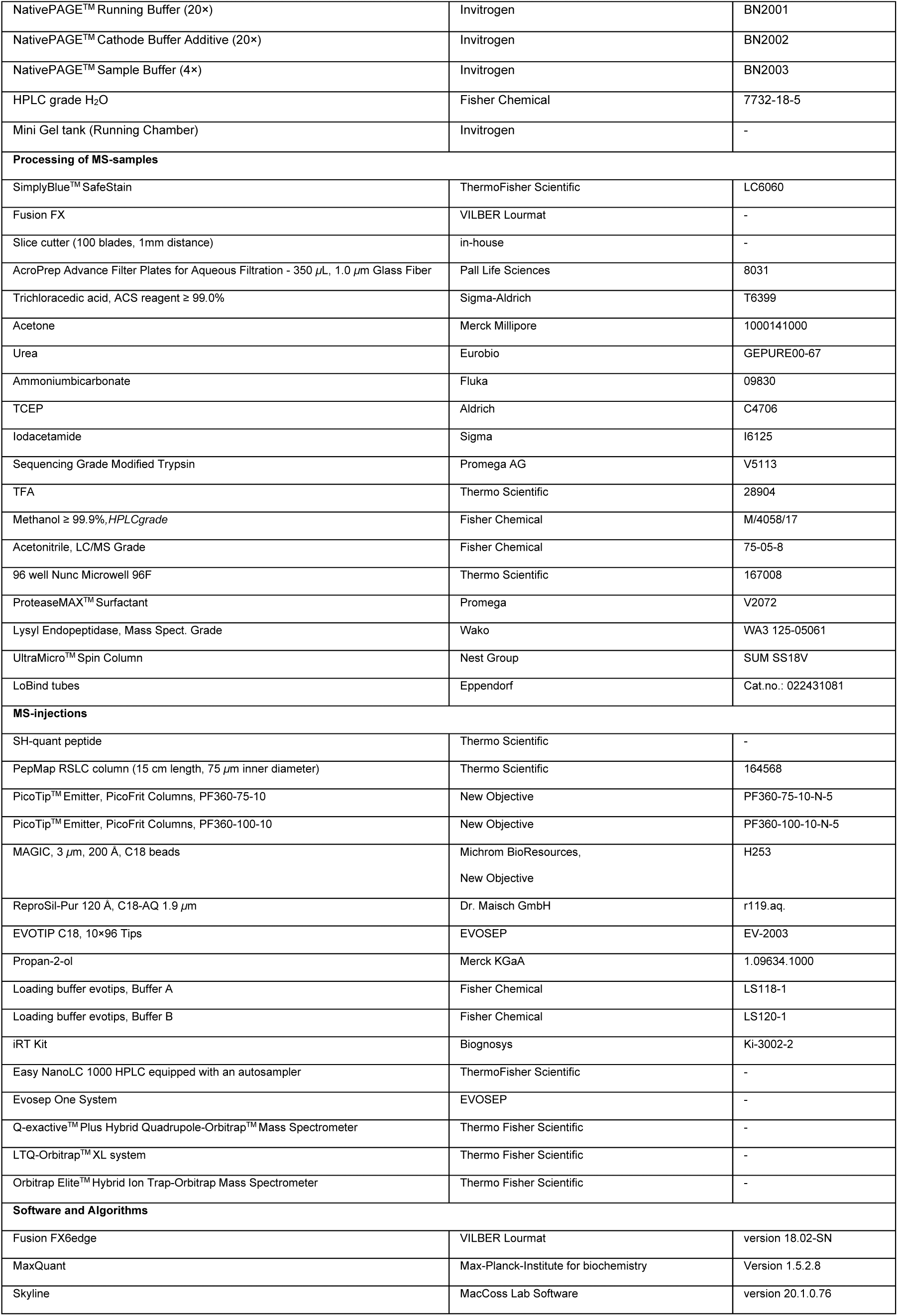
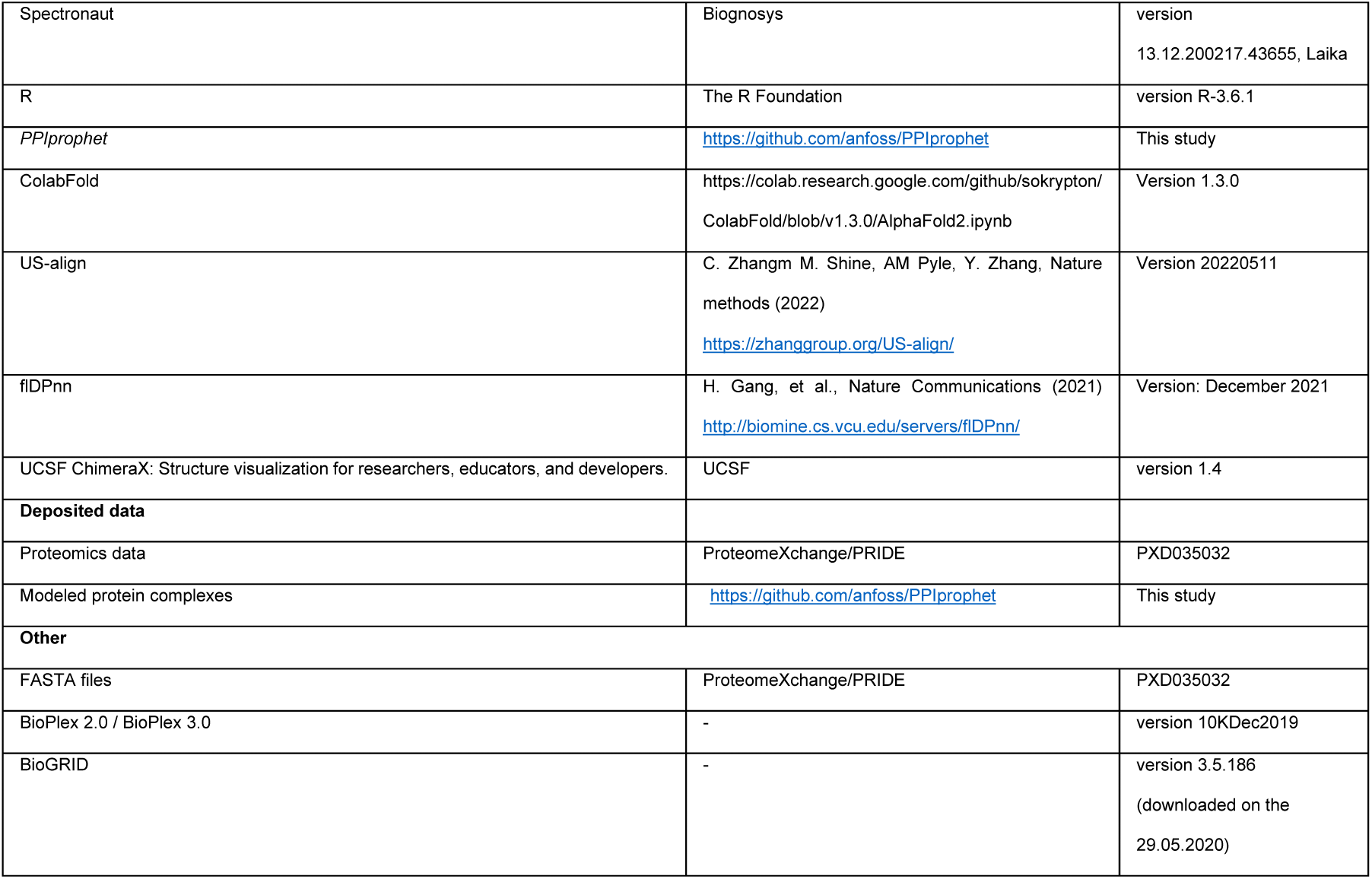

## Extended data figure

**Extended data Fig. 1.**
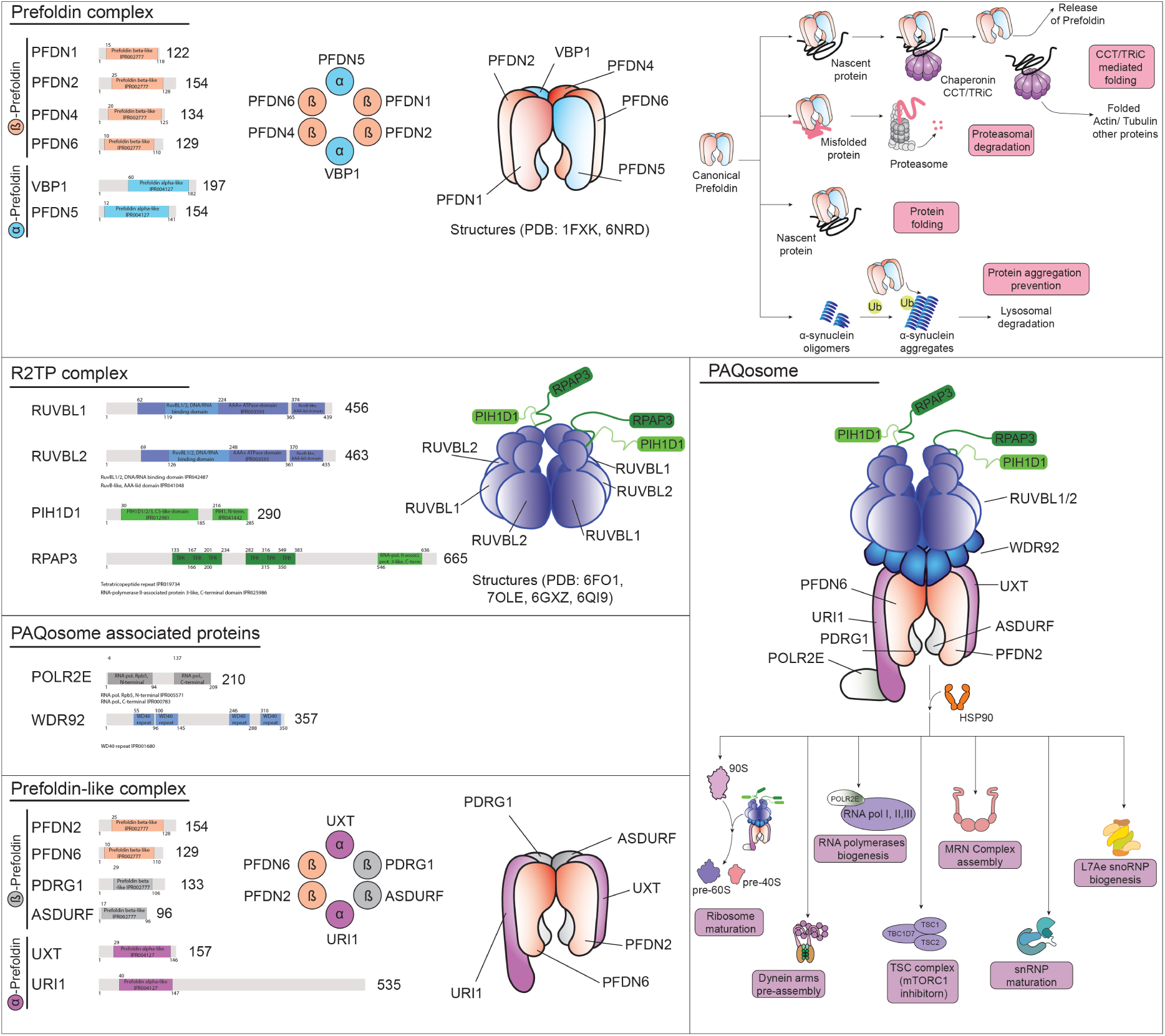
Schematic representation of composition of canonical Prefoldin complex and Prefoldin-like/URI protein complexes. Prefoldin and prefoldin-like protein complexes are reported to be heterohexameric complexes consisting of two α-subunits and four β- subunits. Canonical prefoldin consist of the α-subunits PFDN5 and VBP1 (blue) and the β-subunits PFDN1, PFDN2, PFDN4 and PFDN6 (red). The PFDL complex consists of the shared β-subunits of PFDN2 and PFDN6, the β -subunits of PDRG1, ASDURF (grey) and the α-subunits of UXT URI1 (violet). The R2TP complex, comprises a heterohexameric RUVBL1/RUBL2 ring and one or multiple copies of the adaptor/regulatory subunits RPAP3 and PIH1D1. The R2TP and PFDL-complex are part of the larger PAQosome assembly, which contains two additional subunits, the URI1 associated POLR2E and the WD40 repeat containing protein WDR92, which is likely associated with the R2TP complex. Protein domains were obtained from InterPro. In addition to a sketch of the complexes, the major biological functionalities of canonical Prefoldin and the PAQosome are sketched (see also Lynham et al. 2022 [93] for a more detailed review of the PAQosome and PFD complex).

**Extended data Fig. 2.**
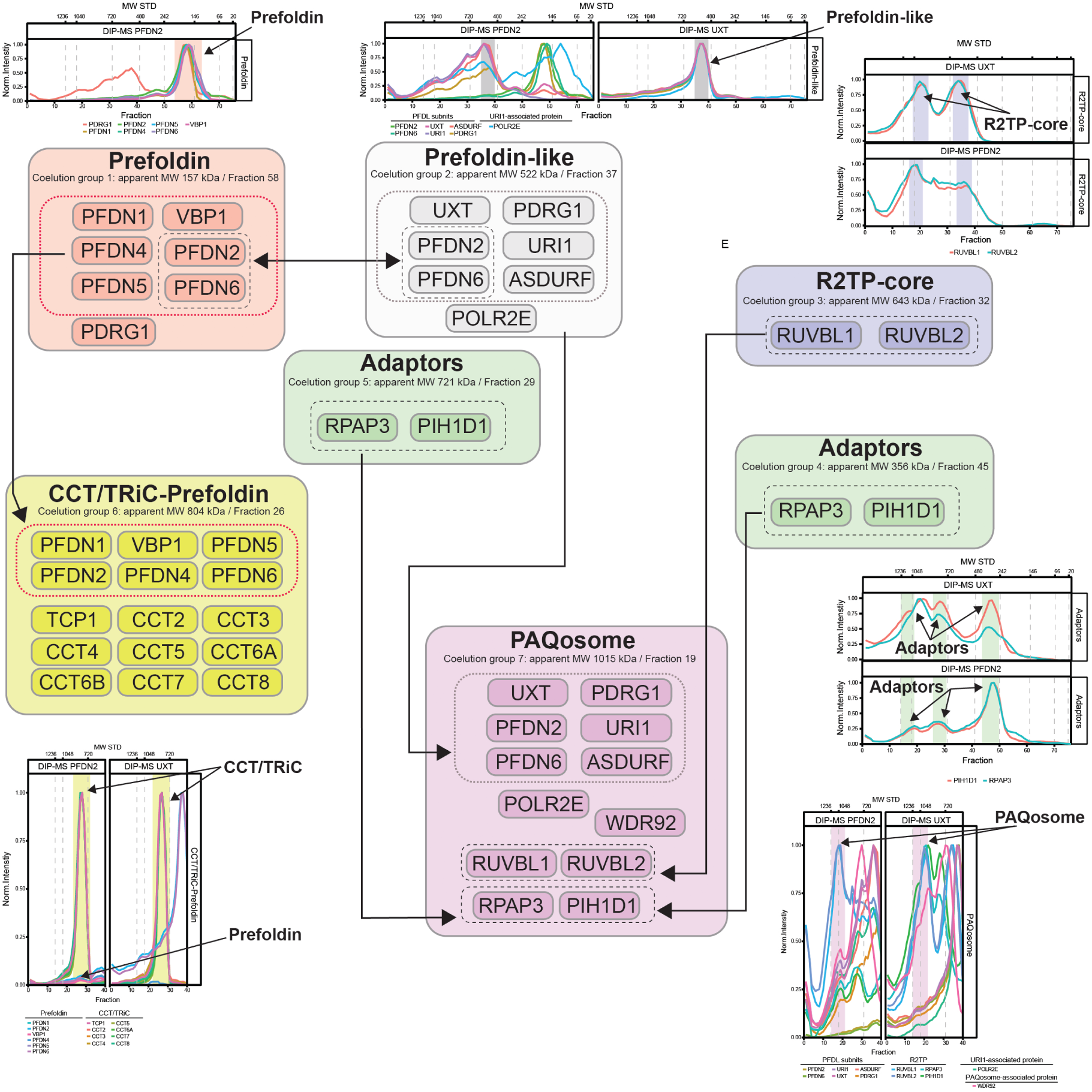
Assembly intermediates and complexes of PFD and PFDL subunits identified across DIP-MS experiments. Across the PFDN2 (n=3) and UXT (n=3) DIP-MS assemblies we characterized 7 distinct coelution peak groups of PFD, PFDL, R2TP and CCT/TRiC core-subunits. The conical PFD complex and PDRG1 eluted at Fraction 58 with an apparent MW of 157 kDa (coelution group 1) and the PFDL-module eluted at Fraction 37 with an apparent MW of 522 kDa (coelution group 2). The R2TP subassembly (coelution group 3 or Fraction 32 apparent MW 643 kDa) and the two adaptor peak subassemblies (apparent MW 356 kDa and 721 kDa or Fraction 45 and 29 assigned as coelution group 4 and 5) were recovered within both DIP-MS experiments. The larger assemblies, CCT/TRiC-Prefoldin (coelution group 6, apparent MW 804 kDa or Fraction 26) and the PAQosome (coelution group 7, apparent MW of 1015 kDa Fraction 19) are indicated at the bottom. Shared subunits and shared complex modules are indicated by arrows and boxes. Below the subassembly, the apparent MW derived from the separation is indicated. Sub-complex stoichiometry was inferred from literature reported assemblies. Apparent MW calibration curve data is in Supplementary Data S16. Within the DIP-MS of UXT the PFD and PFD-CCT/TRiC was not recovered (only residual CCT/TRiC from background). For the coelution groups profile plots, the y-axis represents the average MS2 protein intensities normalized by the maximum protein intensity across the gel slices (x-axis).

**Extended data Fig. 3.**
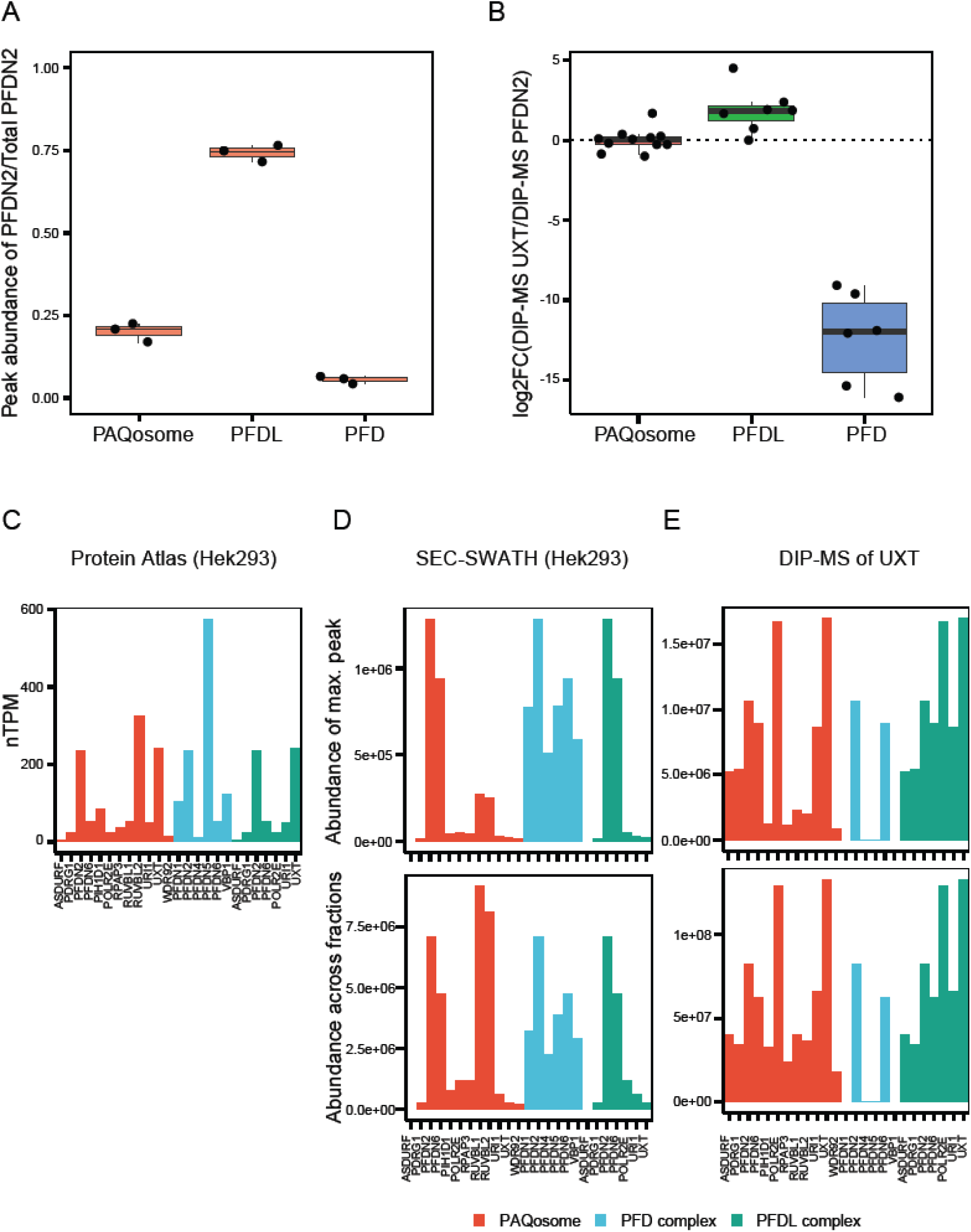
Enrichment of PAQosome, PFDL and PFD in DIP-MS of UXT and PFDN2. **A**. Relative stoichiometry of PFDN2 for the UXT DIP-MS experiment Boxplot showing the ratio of PFDN2 signal in the three major assemblies PFD, PFDL and PAQosome versus the total PFDN2 signal. Box represents the interquantile range and its whiskers 1.5 X IQR. Black line represents median and the replicates are shown with black dots (n=3). **B.** Log2FC of enrichment of PAQosome, PFDL and canonical PFD complexes in the UXT DIP-MS against the PFDN2 DIP-MS. The average across the replicates at the peak group fractions of the complexes was used as proxy for the abundance of the complexes. For each subunit within each peak group the log2FC of the MS2 protein intensity in the UXT against the PFDN2 DIP-MS experiment was calculated. **C.-E.** Abundance comparison of PFD, PFDL and PAQosome core-subunits within transcriptomics, quantitative SEC-MS and DIP-MS. **C.** Normalized transcripts per million (nTPM) of core-subunits for the PAQosome (red), PFD (blue) and PFDL (green). **D.** Peak intensity and summed abundance across all fractions for each subunit in a SEC-SWATH (n=1) experiment and **E.** for DIP-MS of UXT (n=3). For the DIP-MS the average across all three replicates per experiment was considered.

**Extended data Fig. 4.**
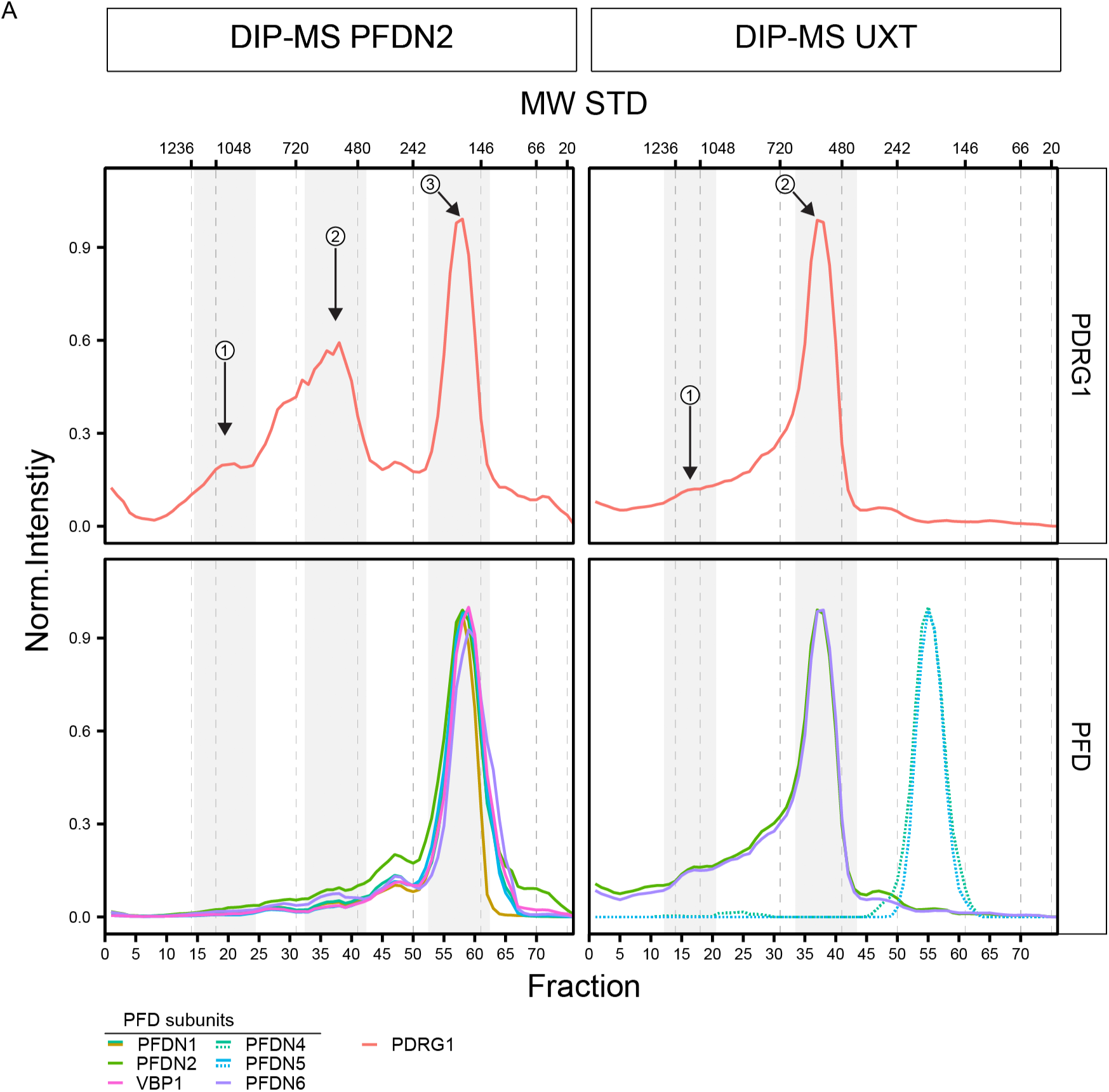
PDRG1 coelution profile in the PFDN2 and UXT DIP-MS experiments. **A**. PDRG1 shows three distinct peaks (labelled 1,2,3) in the DIP-MS experiments of PFDN2, whereas in the UXT DIP-MS only two peaks (1,2) are identified. In both DIP-MS experiments the two peaks at the higher MW belong to the PFDL complex (2) and the PFDL containing PAQosome complex (1). In the PFDN2 DIP-MS the third (3) and most intense PDRG1 peak coelutes with canonical PFD. The intensity shows the smoothed average MS2 protein intensity normalized to the maximum protein intensity across all gel-slices. PFDN4 and PFDN5 coeluting at the PFD peak (dashed lines in the UXT DIP-MS) are contaminant signal in the UXT DIP-MS experiment. The signal is of PFDN4 and PFDN5 is three orders of magnitude lower compared to the PFDN2 signal at the PFDL complex.

**Extended data Fig. 5.**
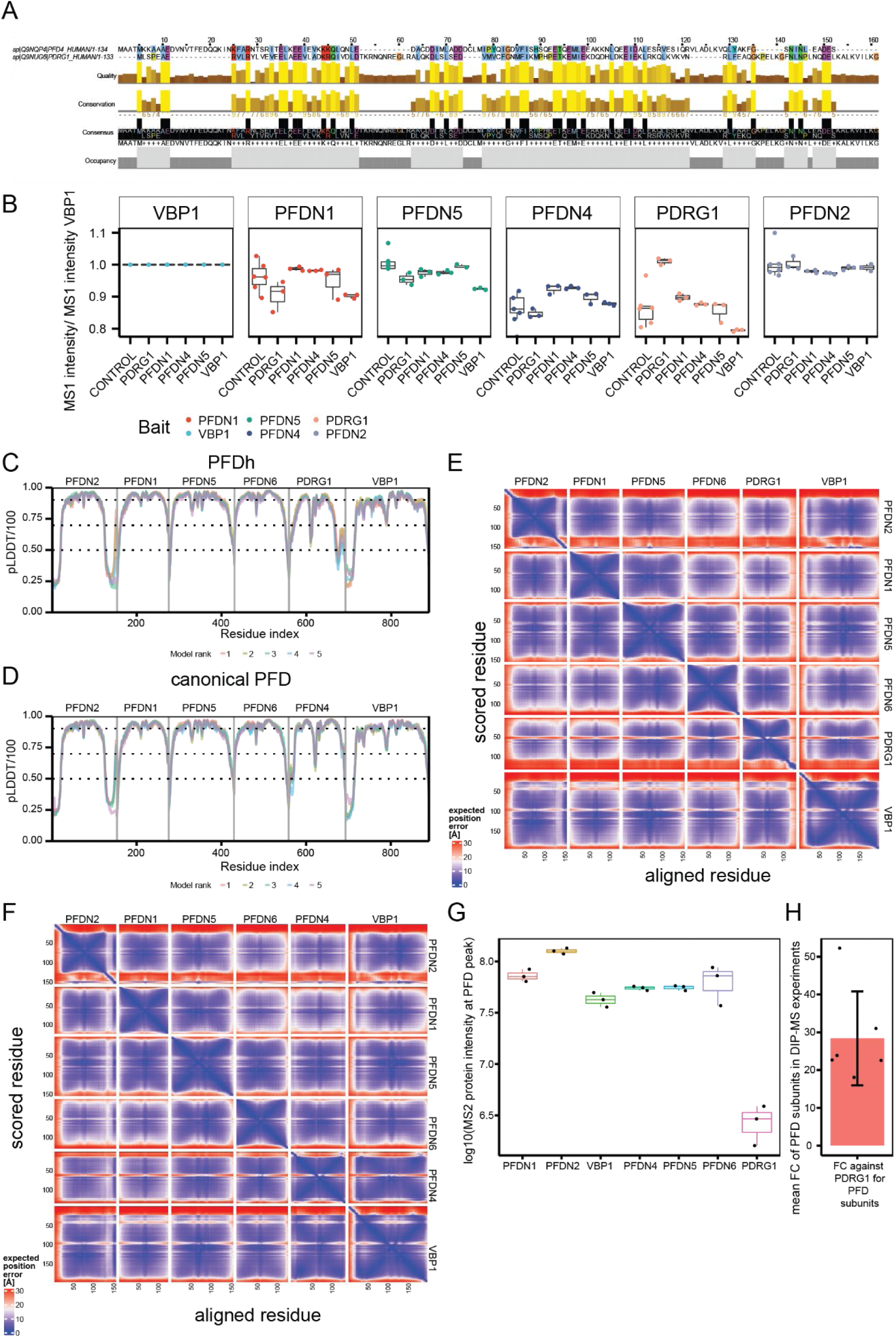
Identification of PDRG1 as subunit of the alternative PFD homolog complex. **A.** Sequence alignment of PDRG1 with PFDN4 indicates large stretches of consensus motives within both proteins, indicating that PFDN4 is the subunit replaced by PDRG1 in the PFD homolog complex. **B.** Ratio to VBP1 comparison on MS1-intensity protein abundance levels in validation AP-MS experiments across different canonical subunits used as baits, showing that PFDN4 ratios were lowest in the AP-MS using PDRG1 as bait, indicative that the recovered PFDN4 interaction with PDRG1 is at the noise level comparable to the level of PDRG1 signal in GFP negative controls. **C.** AlphaFold2 confidence in pLDDT (predicted Local Distance Difference Test) divided by 100 for 5 models of the PFD homolog complex. **D.** AlphaFold2 confidence in pLDDT/100 for 5 models of the canonical PFD complex. **E.** Inter PAE (Predicted Aligned Error) heatmap for the best ranked model of PFDh complex. **F.** Inter PAE (Predicted Aligned Error) heatmap for the best ranked model of the PFD complex. **G.** Comparison of log10 MS2 protein abundance of canonical PFD and PDRG1 in the PFD coelution peak. Points represent signals from the replicates of the DIP-MS experiments. On the right average log2FC against PDRG1, and each point represents the average of a PFD subunits in the PFD-peak across the DIP-MS triplicates.

**Extended data Fig. 6.**
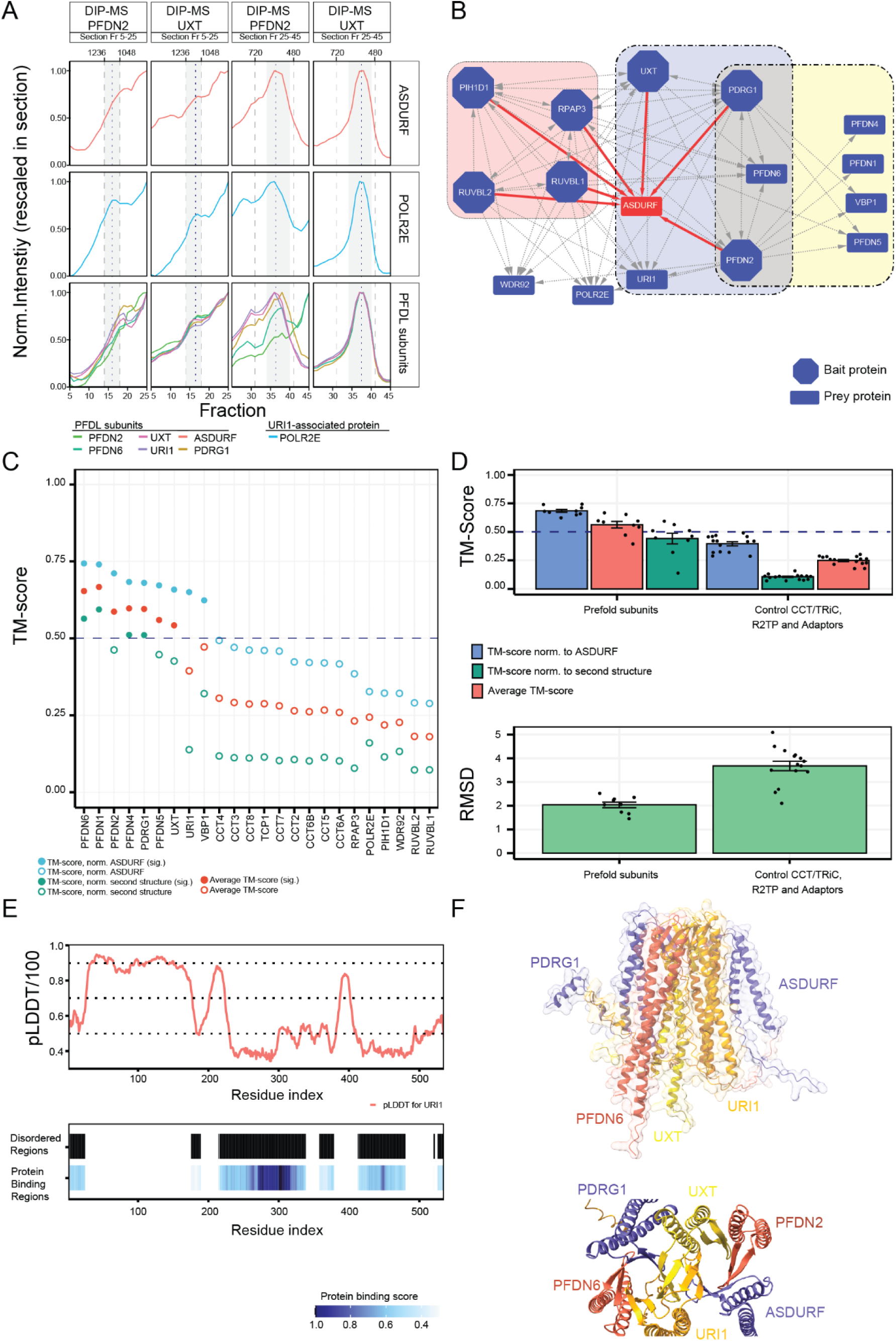
Identification of ASDURF and POLR2E as constitutive PFDL subunits. **A.** Coelution of ASDURF with the PFDL and the PFDL containing PAQosome complex within the PFDN2 and UXT DIP-MS experiments. The MS2 protein intensity was rescaled within the sections (PAQosome section: Fraction 5 – 25 and PFDL section 25 – 45) due to high signal differences between the PFDL and PAQosome complexes. **B.** Interactome derived from AP-MS of core-subunits of the R2TP complex (red background), PFDL (blue background), and canonical prefoldin (yellow background). Interactions are filtered only for high-confidence (Log2FC≥5 and Saint score ≥0.99) and showing only interactions between core-subunits. Baits are depicted as octagons, and red edges represent interactions with ASDURF (shown in red). ASDURF was only recovered with subunits of the PFDL and PFDL containing PAQosome complexes. **C.** Structural alignment of ASDURF AlphaFold model to canonical PFD, PFDL, and CCT/TRiC subunits. The TM-score was either normalized to ASDURF (blue circles) or the second structure (green circles). The mean of the two normalized TM-score is reported (red circles). A significant threshold of 0.5 TM-score was applied (dotted line). Filled circles indicate high structural similarity, empty circles indicate lower to no structural similarity. **D.** Average TM-score and RMSD for Prefoldin subunits and the negative control (subunits of CCT/TRiC, R2TP and adaptor proteins). The average TM-score for prefoldin subunits indicate strong structural similarities between all PFD subunits and ASDURF. The average RMSD of the alignment (2 Å) indicates that the predicted ASDURF model structure shares strong similarities with PFD and PFDL subunits. **E.** AlphaFold2 pLDDT score divided by 100 of URI1 (AF-O94763-F1) plot by residue. Very low pLDDT/100 score (< 0.4) indicate IDRs. At the bottom prediction of IDRs and protein binding regions employing flDPnn [90] indicate large regions IDR within URI1 which explains the poor predictions of these regions in the AlphaFold2 model. **F.** Structural model of the PFDL complex. Subunits are colored. Upper part shows a side view of the complex, the lower part shows a top view, showing the stacked β-sheets.

**Extended data Fig. 7.**
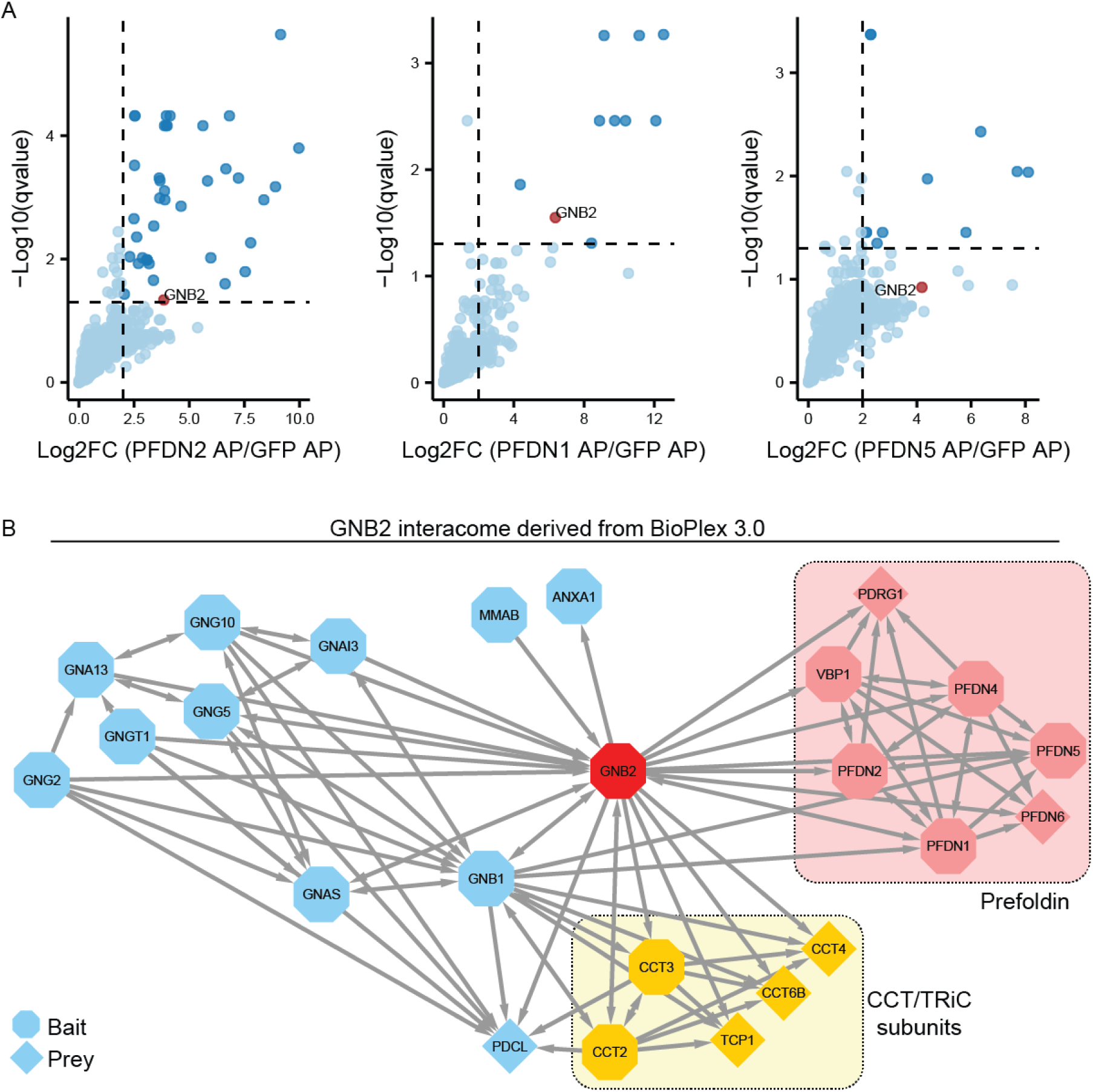
Validation of GNB2 as PFD subunit by reciprocal AP-MS. **A**. Volcano plot for exclusive PFD subunits (PFDN1, PFDN5) and PFDN2, highlighting the recovery of GNB2 as enriched protein versus GFP purification. Dotted lines correspond to a Log2FC=2 and a q-value=5%. **B.** Subnetwork of proteins interacting with GNB2 (red octagon) extracted from BioPlex 3.0 (for HEK293T cells). The reported interactions cover protein-protein interactions between GNB2 and its interactors. The network supports our findings, that GNB2 is interacting with Prefoldin subunits (light red) and CCT/TRiC subunits (yellow). The direction of interaction is indicated as an error on the edges, and the shape of the indicates of the interaction partner was a bait within BioPlex 3.0.

**Extended data Fig. 8.**
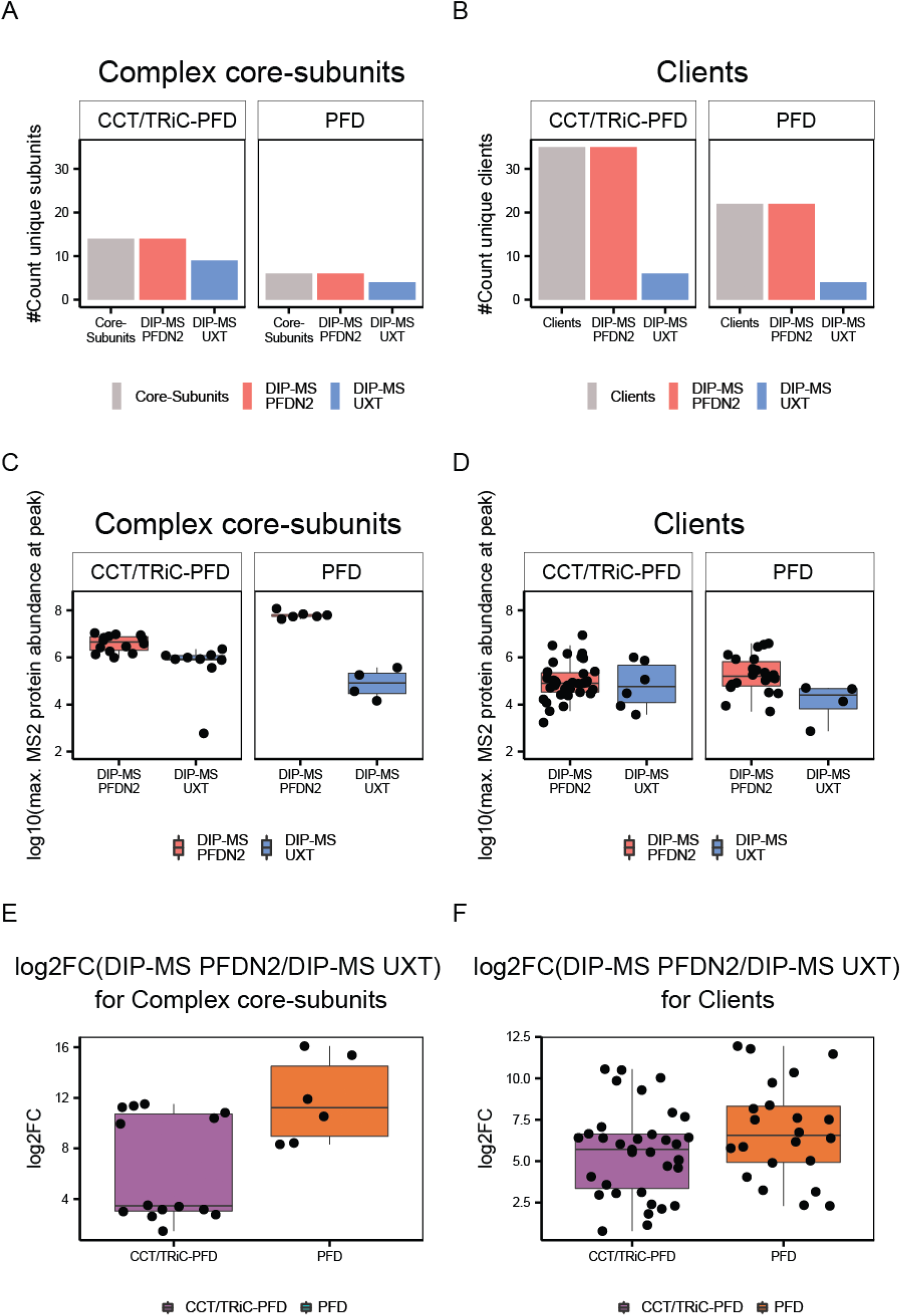
Quantitative comparison of core-subunits and client proteins coeluting with CCT/TRiC-PFD and PFD complexes in the PFDN2 and UXT DIP-MS experiments. A. Recovery of CCT/TRiC-PFD and PFD subunits in the DIP-MS experiments of PFDN2 and UXT. B. Recovery of CCT/TRiC-PFD and PFD clients in the DIP-MS experiments of PFDN2 and UXT. C. Log10 quantitative protein abundance of core-subunits (max. MS2 signal at the CCT/TRiC-PFD or PFD peaks) in the PFDN2 and UXT DIP-MS experiments. The lows abundance of canonical PFD complex in the UXT DIP-MS, indicates that the PFD complex is a background contaminant within this experiment. D. Log10 quantitative protein abundance of clients (max. MS2 signal at the CCT/TRiC-PFD or PFD peaks) in the PFDN2 and UXT DIP-MS experiments. E. Log2FC of core-subunits quantified in the PFDN2 DIP-MS against UXT DIP-MS protein abundance. F. Log2FC of clients quantified in the PFDN2 DIP-MS against UXT DIP-MS on protein abundance.

**Extended data Fig. 9.**
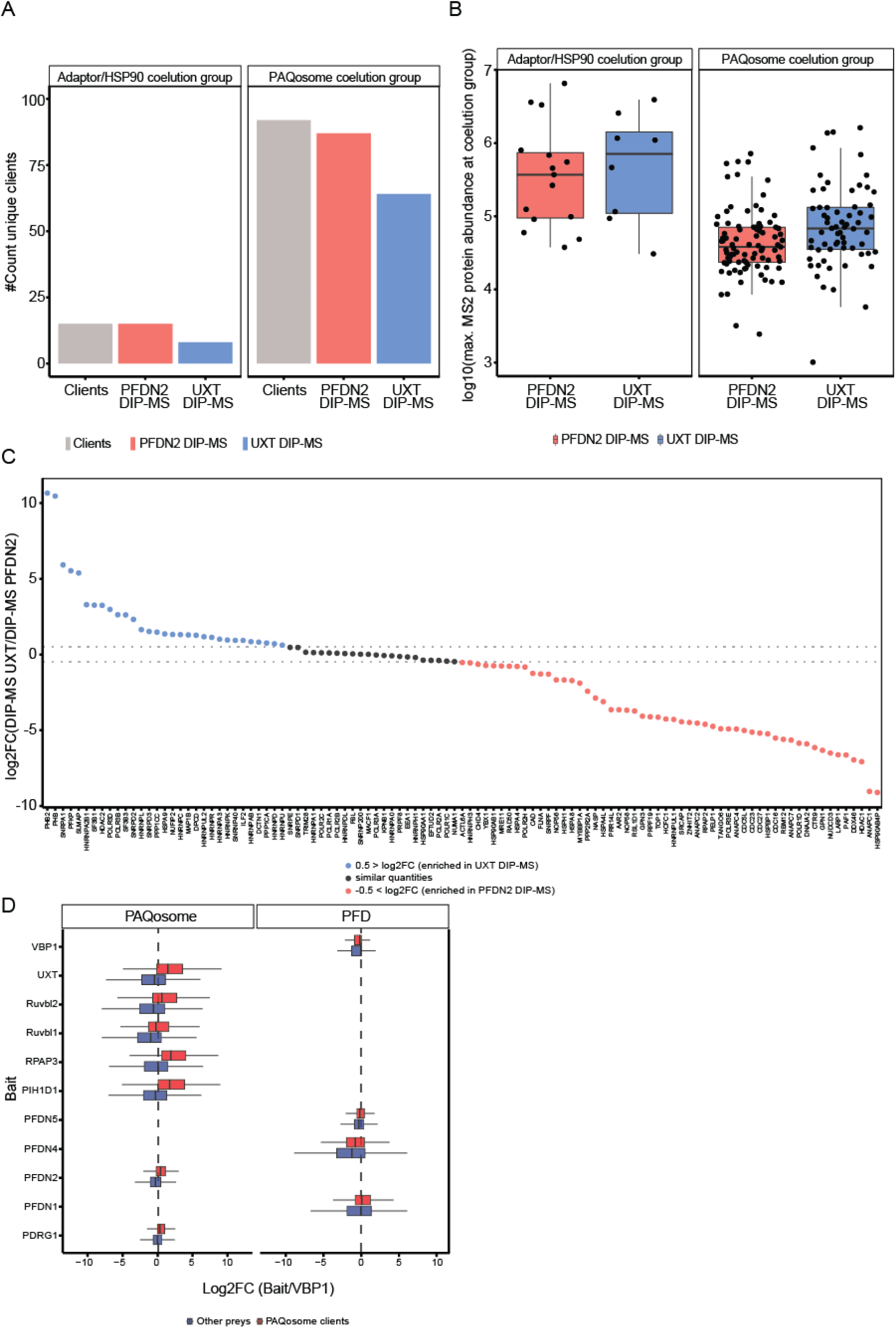
Quantitative comparison of client complex subunits and client proteins coeluting with the PAQosome and the Adaptor/HSP90 assemblies in the PFDN2 and UXT DIP-MS experiments. **A**. Recovery of client complex subunits and client proteins in the DIP-MS experiments, separated by the two coelution groups of Adaptor/HSP90 (red) and PAQosome (blue) coelution groups. The combined number of clients is reported in grey. B. Comparison of client abundance in MS2 protein abundance, averaged maximum intensity of clients across the three DIP-MS experiments for PFDN2 and UXT at the coelution groups of the Adaptor/HSP90 (red) and PAQosome (blue) coelution groups. C. Log2FC of client proteins abundance in UXT DIP-MS divided by the PFDN2 DIP-MS experiment. Missing values were imputed by 1e3 to derive log2FC (indicated in red in the Supplementary Data S.15). Values are ordered from largest to smallest log2FC. Clients recovered with higher signal in UXT DIP-MS (log2FC > 0.5) are reported in blue points, whereas clients quantified higher in the PFDN2 DIP-MS experiment (log2FC < -0.5) are red points. A group of slightly to unchanged protein clients (log2FC < 0.5 and > -0.5) are reported in black points. D. Boxplot showing the enrichment of PAQosome clients (red box) versus the other identified proteins (blue box) across AP-MS. X axis represents the Log2FC calculated across bait proteins (Y axis) versus the corresponding protein abundance in VBP1, used here as representative PFD exclusive subunit. Different columns show PAQosome core components or PFD subunits.

## References

1. Taggart, J.C., et al., Keeping the Proportions of Protein Complex Components in Check. Cell Systems, 2020. 10(2): p. 125–132.

2. Williams, E.G., et al., Systems proteomics of liver mitochondria function. Science, 2016. 352(6291).

3. Hartwell, L.H., et al., From molecular to modular cell biology. Nature, 1999. 402(6761 Suppl): p. C47–52.

4. Gingras, A.C., et al., Analysis of protein complexes using mass spectrometry. Nature Reviews Molecular Cell Biology, 2007. 8(8): p. 645–654.

5. Huttlin, E.L., et al., Dual proteome-scale networks reveal cell-specific remodeling of the human interactome. Cell, 2021. 184(11): p. 3022-+.

6. Hauri, S., et al., A High-Density Map for Navigating the Human Polycomb Complexome. Cell Reports, 2016. 17(2): p. 583–595.

7. Havugimana, P.C., et al., A Census of Human Soluble Protein Complexes. Cell, 2012. 150(5): p. 1068–1081.

8. Kristensen, A.R., J. Gsponer, and L.J. Foster, A high-throughput approach for measuring temporal changes in the interactome. Nat Methods, 2012. 9(9): p. 907–9.

9. Schagger, H. and G. von Jagow, Blue native electrophoresis for isolation of membrane protein complexes in enzymatically active form. Anal Biochem, 1991. 199(2): p. 223–31.

10. Wessels, H.J.C.T., et al., LC-MS/MS as an alternative for SDS-PAGE in blue native analysis of protein complexes. Proteomics, 2009. 9(17): p. 4221–4228.

11. Rudashevskaya, E.L., A. Sickmann, and S. Markoutsa, Global profiling of protein complexes: current approaches and their perspective in biomedical research. Expert Review of Proteomics, 2016. 13(10): p. 951–964.

12. Salas, D., et al., Next-generation Interactomics: Considerations for the Use of Co-elution to Measure Protein Interaction Networks. Molecular & Cellular Proteomics, 2020. 19(1): p. 1–10.

13. Bode, D., et al., Characterization of Two Distinct Nucleosome Remodeling and Deacetylase (NuRD) Complex Assemblies in Embryonic Stem Cells. Molecular & Cellular Proteomics, 2016. 15(3): p. 878–891.

14. Dayebgadoh, G., et al., Biochemical Reduction of the Topology of the Diverse WDR76 Protein Interactome. Journal of Proteome Research, 2019. 18(9): p. 3479–3491.

15. Ciuffa, R., et al., Novel biochemical, structural, and systems insights into inflammatory signaling revealed by contextual interaction proteomics. Proc Natl Acad Sci U S A, 2022. 119(40): p. e2117175119.

16. Heusel, M., et al., Complex-centric proteome profiling by SEC-SWATH-MS. Molecular Systems Biology, 2019. 15(1).

17. Scott, N.E., et al., Interactome disassembly during apoptosis occurs independent of caspase cleavage. Molecular Systems Biology, 2017. 13(1).

18. Vainberg, I.E., et al., Prefoldin, a chaperone that delivers unfolded proteins to cytosolic chaperonin. Cell, 1998. 93(5): p. 863–73.

19. Geissler, S., K. Siegers, and E. Schiebel, A novel protein complex promoting formation of functional alpha- and gamma-tubulin. EMBO J, 1998. 17(4): p. 952–66.

20. Siegers, K., et al., Compartmentation of protein folding in vivo: sequestration of non-native polypeptide by the chaperonin-GimC system. EMBO J, 1999. 18(1): p. 75–84.

21. Martin-Benito, J., et al., Structure of eukaryotic prefoldin and of its complexes with unfolded actin and the cytosolic chaperonin CCT. EMBO J, 2002. 21(23): p. 6377–86.

22. Abe, A., et al., Prefoldin Plays a Role as a Clearance Factor in Preventing Proteasome Inhibitor-induced Protein Aggregation. Journal of Biological Chemistry, 2013. 288(39): p. 27764–27776.

23. Tashiro, E., et al., Prefoldin protects neuronal cells from polyglutamine toxicity by preventing aggregation formation. J Biol Chem, 2013. 288(27): p. 19958–72.

24. Takano, M., et al., Prefoldin prevents aggregation of alpha-synuclein. Brain Res, 2014. 1542: p. 186–94.

25. Comyn, S.A., et al., Prefoldin Promotes Proteasomal Degradation of Cytosolic Proteins with Missense Mutations by Maintaining Substrate Solubility. PLoS Genet, 2016. 12(7): p. e1006184.

26. Djouder, N., et al., S6K1-mediated disassembly of mitochondrial URI/PP1gamma complexes activates a negative feedback program that counters S6K1 survival signaling. Mol Cell, 2007. 28(1): p. 28–40.

27. Mita, P., et al., Analysis of URI Nuclear Interaction with RPB5 and Components of the R2TP/Prefoldin-Like Complex. Plos One, 2013. 8(5).

28. Gstaiger, M., et al., Control of nutrient-sensitive transcription programs by the unconventional prefoldin URI. Science, 2003. 302(5648): p. 1208–1212.

29. Gestaut, D., et al., The Chaperonin TRiC/CCT Associates with Prefoldin through a Conserved Electrostatic Interface Essential for Cellular Proteostasis. Cell, 2019. 177(3): p. 751-+.

30. Boulon, S., et al., The Hsp90 chaperone controls the biogenesis of L7Ae RNPs through conserved machinery. Journal of Cell Biology, 2008. 180(3): p. 579–595.

31. Cloutier, P., et al., R2TP/Prefoldin-like component RUVBL1/RUVBL2 directly interacts with ZNHIT2 to regulate assembly of U5 small nuclear ribonucleoprotein. Nature Communications, 2017. 8.

32. Horejsi, Z., et al., CK2 Phospho-Dependent Binding of R2TP Complex to TEL2 Is Essential for mTOR and SMG1 Stability. Molecular Cell, 2010. 39(6): p. 839–850.

33. Kim, S.G., et al., Metabolic Stress Controls mTORC1 Lysosomal Localization and Dimerization by Regulating the TTT-RUVBL1/2 Complex. Molecular Cell, 2013. 49(1): p. 172–185.

34. Houry, W.A., E. Bertrand, and B. Coulombe, The PAQosome, an R2TP-Based Chaperone for Quaternary Structure Formation. Trends in Biochemical Sciences, 2018. 43(1): p. 4–9.

35. Gillet, L.C., et al., Targeted Data Extraction of the MS/MS Spectra Generated by Data-independent Acquisition: A New Concept for Consistent and Accurate Proteome Analysis. Molecular & Cellular Proteomics, 2012. 11(6).

36. Bekker-Jensen, D.B., et al., Rapid and site-specific deep phosphoproteome profiling by data-independent acquisition without the need for spectral libraries. Nature Communications, 2020. 11(1).

37. Skinnider, M.A. and L.J. Foster, Meta-analysis defines principles for the design and analysis of co-fractionation mass spectrometry experiments. Nature Methods, 2021. 18(7): p. 806-+.

38. Sowa, M.E., et al., Defining the Human Deubiquitinating Enzyme Interaction Landscape. Cell, 2009. 138(2): p. 389–403.

39. Enright, A.J., S. Van Dongen, and C.A. Ouzounis, An efficient algorithm for large-scale detection of protein families. Nucleic Acids Research, 2002. 30(7): p. 1575–1584.

40. Stark, C., et al., BioGRID: a general repository for interaction datasets. Nucleic Acids Research, 2006. 34: p. D535–D539.

41. Kotlyar, M., et al., IID 2018 update: context-specific physical protein-protein interactions in human, model organisms and domesticated species. Nucleic Acids Research, 2019. 47(D1): p. D581–D589.

42. Choi, H., et al., SAINT: probabilistic scoring of affinity purification-mass spectrometry data. Nature Methods, 2011. 8(1): p. 70–U100.

43. Boulon, S., et al., HSP90 and Its R2TP/Prefoldin-like Cochaperone Are Involved in the Cytoplasmic Assembly of RNA Polymerase II. Molecular Cell, 2010. 39(6): p. 912–924.

44. Puri, T., et al., Dodecameric structure and ATPase activity of the human TIP48/TIP49 complex. Journal of Molecular Biology, 2007. 366(1): p. 179–192.

45. Niewiarowski, A., et al., Oligomeric assembly and interactions within the human RuvB-like RuvBL1 and RuvBL2 complexes. Biochemical Journal, 2010. 429: p. 113–125.

46. Gorynia, S., et al., Structural and functional insights into a dodecameric molecular machine - The RuvBL1/RuvBL2 complex. Journal of Structural Biology, 2011. 176(3): p. 279–291.

47. Zhou, C.Y., et al., Regulation of Rvb1/Rvb2 by a Domain within the INO80 Chromatin Remodeling Complex Implicates the Yeast Rvbs as Protein Assembly Chaperones. Cell Reports, 2017. 19(10): p. 2033–2044.

48. Martino, F., et al., RPAP3 provides a flexible scaffold for coupling HSP90 to the human R2TP co-chaperone complex. Nat Commun, 2018. 9(1): p. 1501.

49. Maurizy, C., et al., The RPAP3-Cterminal domain identifies R2TP-like quaternary chaperones. Nature Communications, 2018. 9.

50. Seraphim, T.V., et al., Assembly principles of the human R2TP chaperone complex reveal the presence of R2T and R2P complexes. Structure, 2022. 30(1): p. 156–171 e12.

51. Wepf, A., et al., Quantitative interaction proteomics using mass spectrometry. Nature Methods, 2009. 6(3): p. 203–205.

52. Cloutier, P., et al., Upstream ORF-Encoded ASDURF Is a Novel Prefoldin-like Subunit of the PAQosome. Journal of Proteome Research, 2020. 19(1): p. 18–27.

53. Mirdita, M., et al., ColabFold: making protein folding accessible to all. Nat Methods, 2022. 19(6): p. 679–682.

54. Siegert, R., et al., Structure of the molecular chaperone prefoldin: unique interaction of multiple coiled coil tentacles with unfolded proteins. Cell, 2000. 103(4): p. 621–32.

55. Zhang, C., et al., US-align: Universal Structure Alignments of Proteins, Nucleic Acids, and Macromolecular Complexes. bioRxiv, 2022.

56. Jumper, J., et al., Highly accurate protein structure prediction with AlphaFold. Nature, 2021. 596(7873): p. 583–589.

57. Jeronimo, C., et al., Systematic analysis of the protein interaction network for the human transcription machinery reveals the identity of the 7SK capping enzyme. Molecular Cell, 2007. 27(2): p. 262–274.

58. Cloutier, P., et al., High-resolution mapping of the protein interaction network for the human transcription machinery and affinity purification of RNA polymerase II-associated complexes. Methods, 2009. 48(4): p. 381–386.

59. Melville, M.W., et al., The Hsp70 and TRiC/CCT chaperone systems cooperate in vivo to assemble the von Hippel-Lindau tumor suppressor complex. Molecular and Cellular Biology, 2003. 23(9): p. 3141–3151.

60. Siegers, K., et al., TRiC/CCT cooperates with different upstream chaperones in the folding of distinct protein classes. EMBO J, 2003. 22(19): p. 5230–40.

61. Taipale, M., et al., A quantitative chaperone interaction network reveals the architecture of cellular protein homeostasis pathways. Cell, 2014. 158(2): p. 434–448.

62. Lukov, G.L., et al., Mechanism of assembly of G protein beta gamma subunits by protein kinase CK2-phosphorylated phosducin-like protein and the cytosolic chaperonin complex. Journal of Biological Chemistry, 2006. 281(31): p. 22261–22274.

63. Plimpton, R.L., et al., Structures of the Gbeta-CCT and PhLP1-Gbeta-CCT complexes reveal a mechanism for G-protein beta-subunit folding and Gbetagamma dimer assembly. Proc Natl Acad Sci U S A, 2015. 112(8): p. 2413–8.

64. Kelly, J.J., et al., Snapshots of actin and tubulin folding inside the TRiC chaperonin. Nature Structural & Molecular Biology, 2022. 29(5): p. 420-+.

65. Banks, C.A.S., et al., Differential HDAC1/2 network analysis reveals a role for prefoldin/CCT in HDAC1/2 complex assembly. Scientific Reports, 2018. 8.

66. Pines, A., et al., TRiC controls transcription resumption after UV damage by regulating Cockayne syndrome protein A. Nature Communications, 2018. 9.

67. Mellacheruvu, D., et al., The CRAPome: a contaminant repository for affinity purification-mass spectrometry data. Nature Methods, 2013. 10(8): p. 730-+.

68. Bizarro, J., et al., NUFIP and the HSP90/R2TP chaperone bind the SMN complex and facilitate assembly of U4-specific proteins. Nucleic Acids Res, 2015. 43(18): p. 8973–89.

69. Miron-Garcia, M.C., et al., The prefoldin bud27 mediates the assembly of the eukaryotic RNA polymerases in an rpb5-dependent manner. PLoS Genet, 2013. 9(2): p. e1003297.

70. Kakihara, Y., et al., Nutritional status modulates box C/D snoRNP biogenesis by regulated subcellular relocalization of the R2TP complex. Genome Biology, 2014. 15(7).

71. Lynham, J. and W.A. Houry, The Multiple Functions of the PAQosome: An R2TP- and URI1 Prefoldin-Based Chaperone Complex. Prefoldins: The New Chaperones, 2018. 1106: p. 37–72.

72. Forget, D., et al., The Protein Interaction Network of the Human Transcription Machinery Reveals a Role for the Conserved GTPase RPAP4/GPN1 and Microtubule Assembly in Nuclear Import and Biogenesis of RNA Polymerase II. Molecular & Cellular Proteomics, 2010. 9(12): p. 2827–2839.

73. Carre, C. and R. Shiekhattar, Human GTPases Associate with RNA Polymerase II To Mediate Its Nuclear Import. Molecular and Cellular Biology, 2011. 31(19): p. 3953–3962.

74. Mita, P., et al., URI Regulates KAP1 Phosphorylation and Transcriptional Repression via PP2A Phosphatase in Prostate Cancer Cells. Journal of Biological Chemistry, 2016. 291(49): p. 25516–25528.

75. Go, C.D., et al., A proximity-dependent biotinylation map of a human cell. Nature, 2021. 595(7865): p. 120-+.

76. Henri, J., et al., Deep Structural Analysis of RPAP3 and PIH1D1, Two Components of the HSP90 Co-chaperone R2TP Complex. Structure, 2018. 26(9): p. 1196-+.

77. Hu, L.Z., et al., EPIC: software toolkit for elution profile-based inference of protein complexes. Nature Methods, 2019. 16(8): p. 737-+.

78. Muller, C.S., et al., Cryo-slicing Blue Native-Mass Spectrometry (csBN-MS), a Novel Technology for High Resolution Complexome Profiling. Mol Cell Proteomics, 2016. 15(2): p. 669–81.

79. Pourhaghighi, R., et al., BraInMap Elucidates the Macromolecular Connectivity Landscape of Mammalian Brain. Cell Syst, 2020. 10(4): p. 333–350 e14.

80. Glatter, T., et al., An integrated workflow for charting the human interaction proteome: insights into the PP2A system. Molecular Systems Biology, 2009. 5.

81. Cox, J., et al., Andromeda: A Peptide Search Engine Integrated into the MaxQuant Environment. Journal of Proteome Research, 2011. 10(4): p. 1794–1805.

82. Bateman, A., et al., UniProt: a worldwide hub of protein knowledge. Nucleic Acids Research, 2019. 47(D1): p. D506–D515.

83. Teo, G.C., et al., SAINTexpress: Improvements and additional features in Significance Analysis of INTeractome software. Journal of Proteomics, 2014. 100: p. 37–43.

84. Escher, C., et al., Using iRT, a normalized retention time for more targeted measurement of peptides. Proteomics, 2012. 12(8): p. 1111–1121.

85. Szklarczyk, D., et al., The STRING database in 2017: quality-controlled protein-protein association networks, made broadly accessible. Nucleic Acids Research, 2017. 45(D1): p. D362–D368.

86. Fossati, A., et al., PCprophet: a framework for protein complex prediction and differential analysis using proteomic data. Nature Methods, 2021. 18(5): p. 520-+.

87. Storey, J.D. and R. Tibshirani, Statistical significance for genomewide studies. Proceedings of the National Academy of Sciences of the United States of America, 2003. 100(16): p. 9440–9445.

88. Madeira, F., et al., The EMBL-EBI search and sequence analysis tools APIs in 2019. Nucleic Acids Res, 2019. 47(W1): p. W636–W641.

89. Waterhouse, A.M., et al., Jalview Version 2--a multiple sequence alignment editor and analysis workbench. Bioinformatics, 2009. 25(9): p. 1189–91.

90. Hu, G., et al., flDPnn: Accurate intrinsic disorder prediction with putative propensities of disorder functions. Nat Commun, 2021. 12(1): p. 4438.

91. Evans, R., et al., Protein complex prediction with AlphaFold-Multimer. bioRxiv, 2022.

92. Pettersen, E.F., et al., UCSF ChimeraX: Structure visualization for researchers, educators, and developers. Protein Sci, 2021. 30(1): p. 70–82.

93. Lynham, J. and W.A. Houry, The Role of Hsp90-R2TP in Macromolecular Complex Assembly and Stabilization. Biomolecules, 2022. 12(8).

